# Deciphering Drought Response Mechanisms: Transcriptomic Insights from Drought-Tolerant and Drought-Sensitive Wheat (*Triticum aestivum* L.) Cultivars

**DOI:** 10.1101/2023.10.26.563210

**Authors:** Birsen Cevher-Keskin, Yasemin Yıldızhan, A. Hediye Sekmen-Çetinel, Osman Uğur Sezerman, Buğra Özer, Rümeysa Fayetörbay, Selma Onarıcı, İsmail Türkan, Mahmut Tör

## Abstract

Drought stress poses a significant threat to wheat (*Triticum aestivum* L.) cultivation, necessitating an in-depth understanding of the molecular mechanisms underpinning drought response in both tolerant and sensitive varieties. In this study, 12 diverse bread wheat cultivars were evaluated for their drought stress responses, with particular emphasis on the contrasting performance of cultivars Atay 85 (sensitive), Gerek 79, and Müfitbey (tolerant).

Transcriptomic analysis was performed on the root and leaf tissues of the aforementioned cultivars subjected to 4-hour and 8-hour drought stress and compared with controls. Differentially expressed genes (DEGs) were categorized based on their cellular component, molecular function, and biological function. Notably, there was greater gene expression variability in leaf tissues compared to root tissues. A noticeable trend of decreased gene expression was observed for cellular processes such as protein refolding and cellular metabolic processes like photorespiration as drought stress duration increased (8 hours) in the leaf tissues of drought-tolerant and sensitive cultivars. Metabolic processes related to gene expression were predominantly activated in response to 4-hour and 8-hour drought stress. The drought-tolerant cultivars exhibited increased expression levels of genes related to protein binding, metabolic processes, and cellular functions, indicating their ability to adapt better to drought stress compared to the drought-sensitive cultivar Atay 85. We detected more than 25 differentially expressed TFs in leaf tissues under 4-hour and 8-hour drought stress, while only 4 TFs were identified in the root tissues of sensitive cultivar. In contrast, the tolerant cultivar exhibited more than 80 different TF transcripts in both leaves and roots after 4 hours of drought stress, with this number decreasing to 18 after 8 hours of drought stress. Differentially expressed genes with a focus on metal ion binding, carbohydrate degradation, ABA-related genes, and cell wall-related genes were highlighted. *Ferritin* (*TaFer*), *TaPME42* and *Extensin-like protein* (*TaExLP*), *Germin-like protein* (*TaGLP 9-1*), *Metacaspase-5* (*TaMC5*), *Arogenate Dehydratase 5 (ADT-5)*, *Phosphoglycerate/ bisphosphoglycerate mutase* (*TaPGM*), *Serine/threonine protein phosphatase 2A* (*TaPP2A*), *GIGANTEA* (*TaGI*), *Polyadenylate-binding protein* (*TaRBP45B*) exhibited differential expression by qRT-PCR in root and leaf tissues of tolerant and sensitive bread wheat cultivars.

This study provides valuable insights into the complex molecular mechanisms associated with drought response in wheat, highlighting genes and pathways involved in drought tolerance. Understanding these mechanisms is essential for developing drought-tolerant wheat varieties, enhancing agricultural sustainability, and addressing the challenges posed by water scarcity.

## Introduction

Bread wheat, *Triticum aestivum* L. is one of the staple crops for many countries. According to the Food and Agriculture Organization of the United Nations (FAO), wheat production has been estimated to be 766.5 million tons in 2020 [1] and the requirement for wheat is expected to rise by 60% by 2050. Drought is a major issue affecting grain yield, kernel weight, and end-use quality at the heading and grain filling stages of wheat [2]. This is particularly problematic factor for wheat agriculture in arid regions, including the central and eastern Anatolian regions of Turkey. Yield losses could reach up to 80% in some years, especially in central Turkey, where groundwater resources have been nearly depleted due to the excessive use for irrigation, further exacerbating the problem [3]. Flowering and grain development stages are the most sensitive to drought stress, which causes decreases in the yield and grain protein quality. In addition, with the effects of climate change, wheat production might go down by 29% [4]. These predictions clearly show that the improvement of drought tolerance in wheat is of great significance for the global food security in the near future. Genetic studies and new approaches to improve wheat productivity under drought conditions is an urgent priority [5].

Drought stress tolerance is a complex trait that involves physiological, biochemical, and molecular mechanisms. Several mechanisms enabling adaptation to drought stress have been identified in drought-tolerant plants, including the reduction of water loss by improving stomatal resistance, the increase of water uptake by developing large and deep root systems, and the accumulation of osmolytes such as proline, glycine-betaine, sugars (mannitol, sorbitol, and trehalose), and glutamate have been identified in drought-tolerant plants to adapt to drought stress [6, 7, 8].

Plant responses to drought stress start with the stimulation of signal transduction cascades. The activation of several transcription factors and regulators initiates the induction of several molecular and cellular mechanisms. Depending on the genetic background, the response to drought stress varies considerably. Moreover, inter- and intra-species changes in drought resistance are also known [9].

A number of transcriptome and proteome profiling and genetic manipulation studies have identified several genes such as *Zeaxanthin epoxidase* (*ZEP*), *9-cis-Epoxycarotenoid dioxygenase* (*NCED*), *Serine/threonine protein kinase* (*SnRK2*), *Dehydration-responsive element binding factor 1* (*DREB1B*) and plasma membrane intrinsic proteins genes (*PIPs*) with potential roles in drought tolerance mechanisms [10, 11, 12, 13, 14, 15, 16].

Microarray and RNA-seq analysis have detected abiotic stress response genes, especially those involved in response to drought stress in different plants [17, 18]. In contrast to microarray methods, sequence-based RNA-seq analysis determines the cDNA sequence. For this reason, RNA-seq offers a far more precise measurement of transcript levels and their isoforms than the other methods. Photosystem components, carbohydrate metabolism, antioxidant enzymes, and tricarboxylic acid cycle related genes have been identified as being responsible for drought tolerance in wheat [19]. During the reproductive stages, over 300 differentially expressed genes related to many significant processes, such as photosynthetic activity, stomatal movement, and floral development have been identified in wheat under drought stress [20]. Several types of transcription factors, such as WRKY, ERF, NAC, bHLH, bZIP, HD-ZIP, dehydrins, heat shock proteins, proteinase inhibitors, and glutathione transferase, have been identified as the main differentially expressed genes in wheat under drought conditions [21].

Genes encoding glutathione S-transferase (GST), RAB, rubisco, helicase, and vacuolar acid invertase are known to be drought-related genes, and their expression is affected by drought stress in different species [22, 23, 24, 25]. Late embryo abundant (LEA) proteins accumulate under stress conditions such as drought, salinity, and low temperatures. Expression profile analysis determined that most of the LEA genes were expressed at a higher rate in drought-resistant varieties than in sensitive ones [26]. The accumulation of members of the DHN family has been linked to stress tolerance involving dehydration in several species, including sunflower [27], barley [26], and wheat [28].

Bogard et al. (2021) showed that genotypic characteristics related to abiotic stress tolerance should be taken into account in the selection of suitable wheat for breeding in different regions [29]. Those authors developed a marker-based statistical model has been developed for the prediction of phenology parameters in wheat and simulated genotype stress avoidance frequencies of frost and heat stress at different locations; the model’s predictions were validated by observing grain yields in a real trial network have been evaluated in low frost and heat risk periods at each location [29]. Since the drought stress relation of some of the genes have not been completely identified yet, our knowledge of genes involved in drought response is still incomplete.

This study is aimed to the discovery of genes that are responsive to drought stress in bread wheat (*Triticum aestivum* L.). Through physiological screening, we discerned wheat cultivars displaying varying levels of sensitivity and tolerance to drought. Leveraging RNA-Seq technology, we probed the expression profiles of drought-responsive genes within the leaves and roots of three distinct wheat cultivars following exposure to different drought stress conditions. Our investigation unveiled a considerable number of genes exhibiting either elevated or decreased level of expression in both drought-tolerant and sensitive bread wheat cultivars. Subsequently, select differentially expressed genes (DEGs) were validated using quantitative real-time polymerase chain reaction (qRT-PCR). The insights gained from this research have the potential to inform the development of drought-tolerant wheat varieties, employing diverse methodologies, including genome editing techniques.

## Materials and Methods

### Plant Growth and Water Stress Treatment

Twelve *T. aestivum* cultivars originating from Turkey were selected as the most promising drought-stress-tolerant and sensitive cultivars (Supplementary Table S1). The seeds were surface sterilized (5 min with 10% EtOH and 5 min with 5% hypochlorite) and pre-germinated in Petri dishes for 10 days on wet filter paper at 4°C in the dark. Seedlings were grown in 1.5 L plastic pots containing a turf: soil: sand (3: 3: 1) mixture at 18-20°C with 60–70% relative humidity in a controlled growth room. Seedlings of a similar germination stage were transferred to pots, and for each cultivar, three pots were used for control and three for the drought stress.

### Drought Stress Treatment

The drought stress treatment (progressive drought stress) was started 3 weeks after transferring the seedlings to the pots and carried out by withholding water from the stress treated pots.

A regular watering regime was carried out for the control plants every day. Soil Water Content (SWC) measurements were taken during the stress. At the end of the tenth day of drought treatment, Relative Water Content (RWC) measurements were calculated for each cultivar as described [30]. All plants were harvested at the end of the 10^th^ day of drought treatment. Harvested tissues were directly frozen in liquid nitrogen and stored at -80°C till use. For each pot, three different measurements were taken in the afternoon for every day [31]. Based on the physiological data (RWC, SWC), from the three biological replicates of each cultivar, drought-sensitive and drought-tolerant bread wheat cultivars were identified. The cultivars Gerek 79 and Müfitbey were selected as drought tolerant and Atay 85 was selected as drought-sensitive for use in further subsequent transcriptomal profiling experiments (Supplementary Figure S1).

### Soil Water Content

The Time Domain Reflectometry (TDR) Soil Moisture System (Spectrum Technologies, Illinois) was used for the estimation of the mean soil moisture. During the progressive drought stress application, soil moisture ratios were measured in pots of drought and control plant samples for each of the 12 cultivars every day.

### The Relative Water Content

At the end of 10 days of drought stress, leaf tissues (the third youngest leaf) were collected for RWC measurements. RWC quantifications were performed as described by Barr and Weatherley (1962) [30]. Fresh leaves (0.5 g) were cut into 1-cm-long fragments and weighed for their fresh weight (FW), then saturated in water for 8 h at 4 ^0^C and weighed for their turgid weight (TW). Subsequently, the samples were dried in an oven at 80 ^0^C for 24 h, and the dry weight (DW) was measured. The RWC was calculated by using the formula (FW-DW)/(TW-DW) X 100%. (Supplementary Figure S2).

### Shock Dehydration Stress

To identify more rapid changes in drought related gene expression, shock dehydration stress (4h and 8h) was carried out with the drought-tolerant cultivars (Müfitbey and Gerek 79) and drought-sensitive cultivar (Atay 85). Seeds were surface sterilized in 70% EtOH for 5 minutes and in 30% sodium hypochlorite for 10 minutes. Subsequently, seeds were rinsed six times with sterile distilled water for 2 minutes and pre-germinated in Petri dishes for 10 days at 4°C in the dark. After germination, seedlings were transferred to 10 L plastic pots containing moistened perlite for growth. Seedlings of a similar developmental stage were transferred to a continuously aerated ½ Hoagland’s solution renewed every 3 days, and grown under controlled conditions (16h photoperiod, temperature 22/18°C and relative humidity 60%). Shock dehydration stress was applied to Gerek 79, Atay 85 and Müfitbey cultivars by removing them from hydroponic culture and keeping them on the bench for 4 and 8 hours at RT. Control samples were not removed from the hydroponic culture during this period and were harvested at the 4th and 8th hours without exposing them to stress (Supplementary Figure S1).

### Isolation of Total RNA

Total RNA isolation was performed from leaf and root tissues using the RNeasy Plant Mini kit (Qiagen, Hilden, Germany) according to the manufacturers’ instructions. RNAse-free DNaseI (Roche Applied Science GmbH, Germany) digestion and purification were carried out for the elimination of the genomic DNA from total RNA as described [31]. Purified RNA quality was evaluated using a Bioanalyzer (Agilent, USA) and only those samples with RIN (RNA integrity number) scores of 8.0 and greater were used in RNAseq analysis.

### RNA Sequencing

Two tissues (leaf and root) of the three biological replicates of each cultivarwere analyzed for each condition (4 h and 4 h Control; 8 h drought and 8 h Control), and two tissues (leaf and root), resulting in a total of 72 samples (3 genotypes × 4 conditions × 3 replicates x 2 tissues). The RNAseq library for each sample was prepared with a 1250 ng of total RNA using the TruSeq RNA Sample Preparation kit (Illumina) according to the manufacturer’s instructions. Paired-end sequencing was performed with a current next generation sequencing instrument, HiSeq2000 (Illumina, user guide; Part# 15011190 Rev. H) using TruSeq SBS Kit v3 (cBot-HS) (Illumina, user guide; Part#15023333 Rev. B). The prepared libraries were enriched using 15 cycles of PCR and purified by the QIAquick PCR purification kit (Qiagen). The Agilent 2100 Bioanalyzer was used to control the size and purity of the samples using the Agilent High Sensitivity DNA Kit. A total of 12 indexes were prepared for 72 samples and run on Illumina HiSeq 2000 for 6 lanes. The enriched libraries were diluted with the elution buffer to a final concentration of 10 nM. Sequencing was performed on each library to generate 100-bp PE reads for transcriptome sequencing on an Illumina High-Seq 2000 platform.

### Differential Gene Expression Analysis

The quality control was performed for the Illumina paired-end sequencing files of each sample. FastQC Software” was used for the detection of faulty sequences [32]. RNA-seq data were trimmed using the Fastx Toolkit (http://hannonlab.cshl.edu/fastx_toolkit) [33]. After quality control, *de novo* assembly was carried out from a total of 311 GB of transcript data. The assembly was performed as recommended by Duan et al. (2012) [34]. The resultant data were evaluated using the software “Trinity Assembly”, which combines three independent software modules (Inchworm, Chrysalis and Butterfly) and 323 Mbs of FASTA files were obtained. To remove the expected redundancy in this assembly file, “the cd-hit-est tool” to place the contigs into clusters was applied, so that a sequence is not represented more than once in our reference assembly. Subsequently, the RNA-seq data were mapped to our *de novo* reference genome using Bowtie (https://bowtie-bio.sourceforge.net/index.shtml). The resulting mapped reads were evaluated by using the RSEM tool to obtain Fragments per Kilobase of transcript per Million mapped reads (FPKM) data. FPKM files belonging to each sample were subjected to pairwise comparison using the edgeR differential expression tool, which is included in the R-Bioconductor package [35]. Through differential expression analysis, we pooled replicates belonging to each condition into a single file by averaging the counting information corresponding to each gene. As a result, comparisons between different conditions were carried out and differentially expressed transcripts were obtained. However, some transcripts were not informative, as they were not annotated due to a lack of well-annotated reference genome. In this case, the Trinotate annotation tool (https://rnabio.org/module-07-trinotate/0007/02/01/Trinotate/) [36] was used which uses various well referenced methods for functional annotation including homology search for known sequence data (NCBI-BLAST), protein domain identification (HMMER/PFAM), protein signal prediction (singalP/tmHMM), and comparison to currently curated annotation databases (EMBL Uniprot eggNOG/GO Pathways databases) have been applied. Functional enrichment terms were filtered by a given threshold, False Discovery Rate (FDR) ≤ 0.05. We took the negative logarithm of base 2 of Fold Change (FC) values of the corresponding enrichment terms. The color intensity, based on the adjusted logarithmic scale of FC values, demonstrates the level of significance of each term. If there was no log2FC score for the corresponding enriched term, this was depicted as white in the heatmap.

### Primer Design for qRT-PCR

Primers were designed for the selected genes using FastPCR and Primer 3 programs. The quality of the primers was validated by BLASTn queries against the entire wheat EST unigene set. The primers, wherever possible, were designed spanning an intron or intron-intron junctions to detect any genomic DNA contamination. All the primers were adjusted to 100-140 bp amplicon size and 55 °C annealing temperature and controlled by conventional PCR by housekeeping genes (β actin, EF-1 and EF2 primers).

### cDNA Synthesis and qRT-PCR

First-strand cDNA was synthesized by reverse transcribing 1 μg of total RNA in a final reaction volume of 20 μl using MMLV reverse transcriptase (Roche High Fidelity cDNA synthesis kit) according to the manufacturer’s instructions. All the cDNA samples were controlled by conventional PCR with housekeeping genes (beta actin, EF1 and EF2) primers. Differentially expressed transcripts were analyzed with SYBR Green Mix (Roche FastStart Universal SYBR Green Master) and specific primers (Supplementary Table S2). Experimental design was performed by IQ5 System (BioRad Laboratories, Hercules, USA) as described by Cevher-Keskin et al. (2011).[37] Three technical replicates were used for each experiment to quantify the transcript level accurately. The relative abundance levels of all gene specific transcripts for different reactions were normalized with respect to the loading standard, housekeeping gene. The relative fold expression differences were calculated using the comparative CT method [38]. Finally, the ΔCT values for all transcripts were averaged across all treatments and experimental replicates. The gene expression was normalized by using EF-α1 and EF-α2 as a housekeeping gene. Error bars are the standard deviation of qRT-PCRs each performed in triplicate. Normalized expression (ΔΔCq) analysis mode was used for each analysis.

### Accession numbers

The datasets generated in the current study are available in Sequence Read Archive (SRA) under accession numbers SRR25998966, SRR25998965, SRR25998964, SRR25998974, SRR25998971, SRR25998968, SRR25998986, SRR25998983, SRR25998980, SRR25998977, SRR25998984, SRR25998981, SRR25998978, SRR25998975, SRR25998972, SRR25998969, SRR25998963, SRR25998985, SRR25998982, SRR25998979, SRR25998976, SRR25998973, SRR25998970, and SRR25998967 and are accessible via BioProject accession.

## RESULTS

### Selection of the Drought-Tolerant and Drought-Sensitive Cultivars

In the present study, 12 bread wheat (*Triticum aestivum* L.) cultivars with diverse genetic backgrounds were used for the selection of the most promising drought stress tolerant and sensitive cultivars (Supplementary Figure S1). Soil Water Content (SWC) measurements were taken during the drought stress induction. Althought the RWC decreased during the drought experiment. by the end of the 10^th^ day of drought treatment, cv. Atay 85 showed a significant decrease of RWC compared to the other varieties, and cvs. Gerek 79 and Müfitbey showed the least decrease. The RWC levels of the sensitive variety Atay 85 was identified as lower than 70% in (Supplementary Figure S2). Based on these results, cvs. Gerek 79 and Müfitbey were selected as drought tolerant whilst cv. Atay 85 was selected as drought-sensitive.

### Identification of DEGs

RNA-seq analysis was carried out on the root and leaf tissues of selected varieties Gerek 79, Müfitbey and Atay 85 subjected to 4h or 8h drought-stress shock or no stress to reveal the differences in the transcript levels (Supplementary Figure S3). Genes tat were differentially expressed genes in the root and leaf tissues of drought stressed bread wheat cultivars compared to the controls were classified according to their biological process, cellular component, and molecular function by the agriGO program in root and leaf tissues [39] (Supplementary Figures S4-S9). The distribution of the expression levels of the genes in the leaf tissues was more variable than that in the root tissues. In Atay 85, 8h drought treated root tissues with a 0.01 threshold, genes with a differentially increased expression fell into the following categories: GO:0015078∼hydrogen ion transmembrane transporter activity, GO:0015077∼monovalent inorganic cation transmembrane transporter activity GO:0022890∼inorganic cation transmembrane transporter activity related genes. In contrast, in tolerant cultivar Müfitbey, genes in the categories GO:0034404∼nucleobase, nucleoside and nucleotide biosynthetic process, GO:0034654∼nucleobase, nucleoside, nucleotide, and nucleic acid biosynthetic process, GO:0016469∼proton-transporting two-sector ATPase complex, GO:0045259∼proton-transporting ATP synthase complex, GO:0044271∼nitrogen compound biosynthetic processes showed increased expression level. In Gerek 79 leaf tissues of the 8h drought stress treatment, genes in the GO:0005506∼iron ion binding, GO:0046906∼tetrapyrrole binding, GO:0009767∼photosynthetic electron transport chain related categories were increased in expression (Figure 1, 2, and 3).

**Figure 1.**
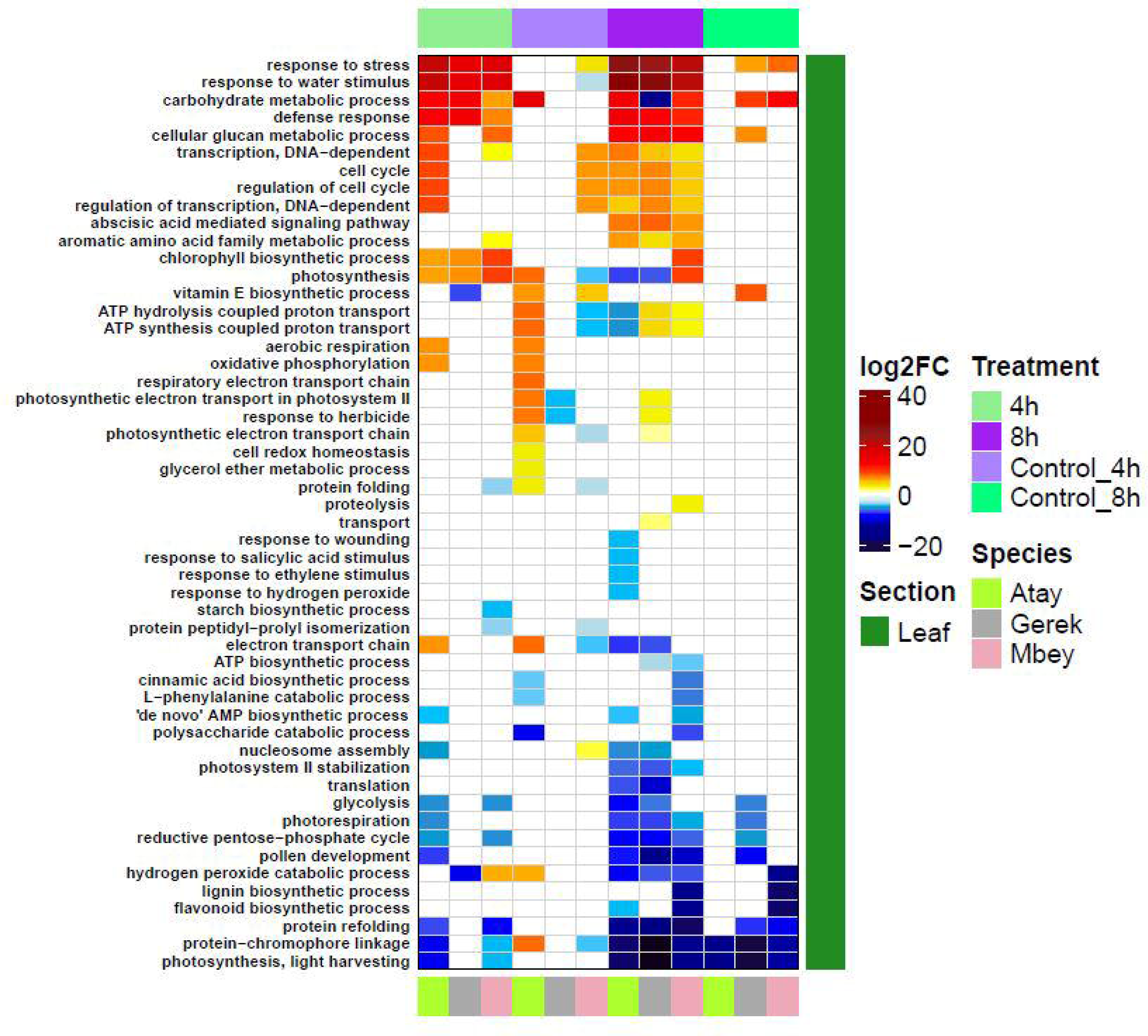
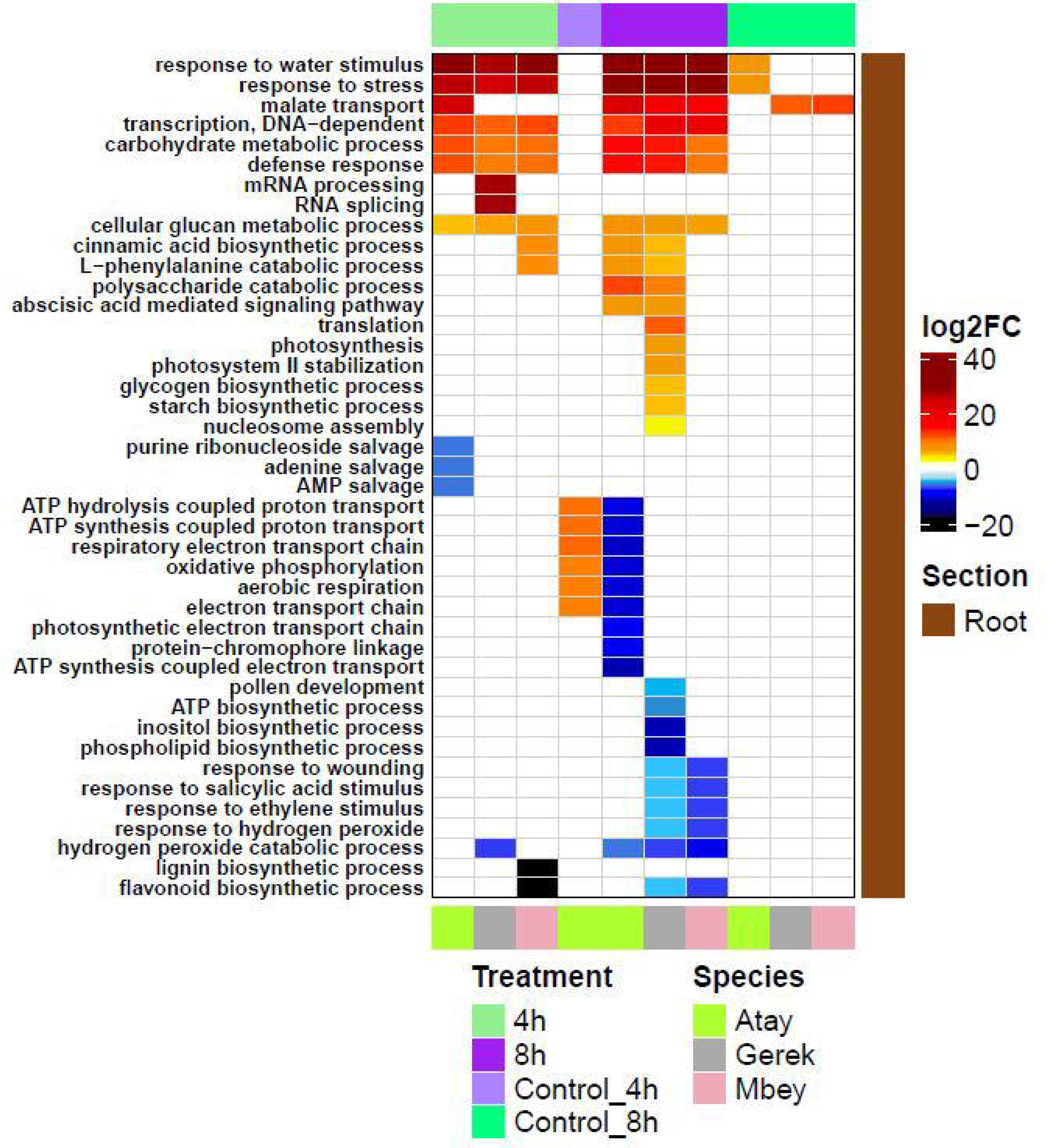
Analysis of biological functions under 4h and 8h drought stress and control groups in leaf and root tissues of the drought-tolerant (Gerek 79, Müfitbey) and sensitive (Atay 85) cultivars. Red and blue colours show the higher and lower expression values, respectively, where Atay 85 is assigned in green, Gerek 79 in grey, and Müfitbey in pink.

**Figure 2.**
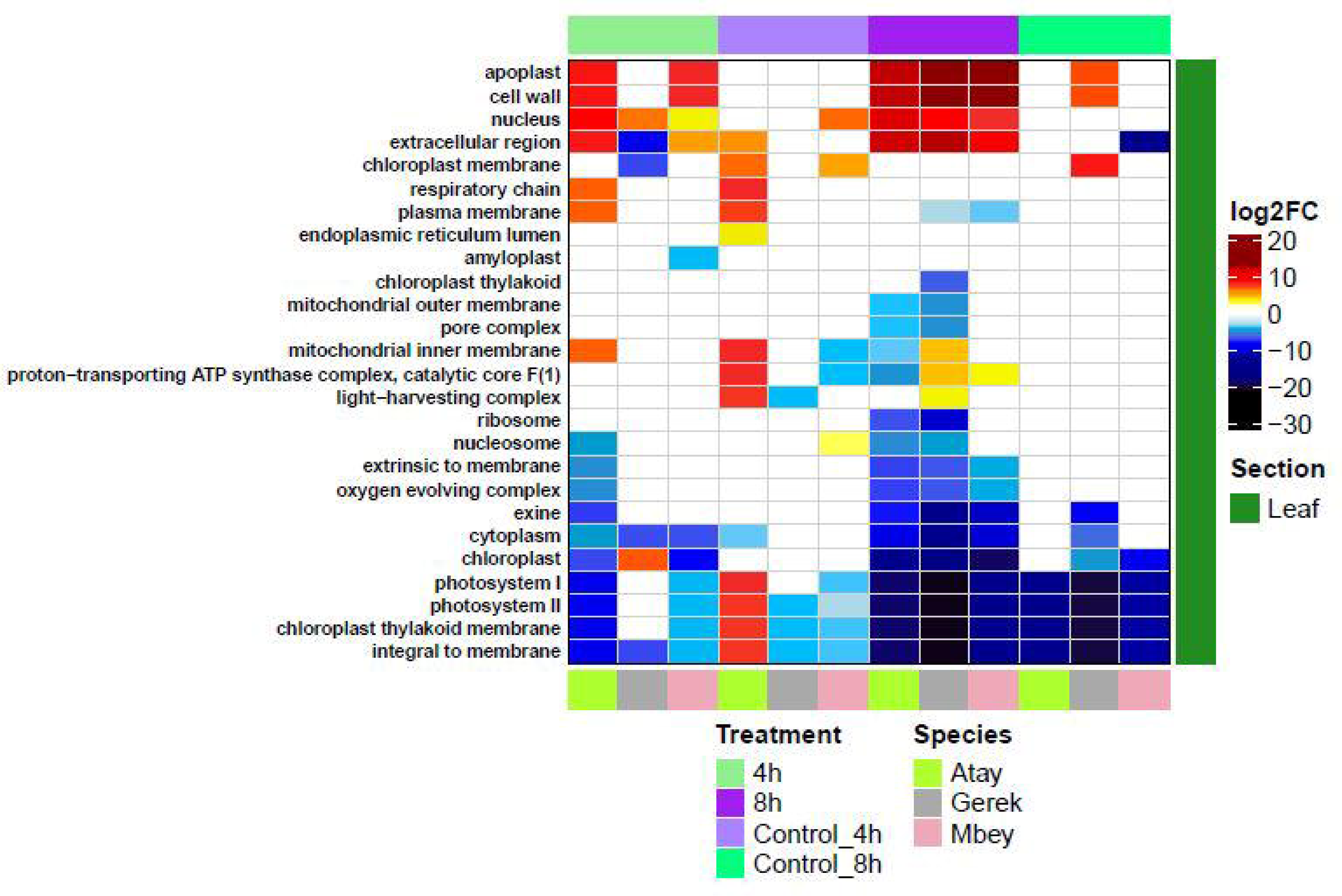
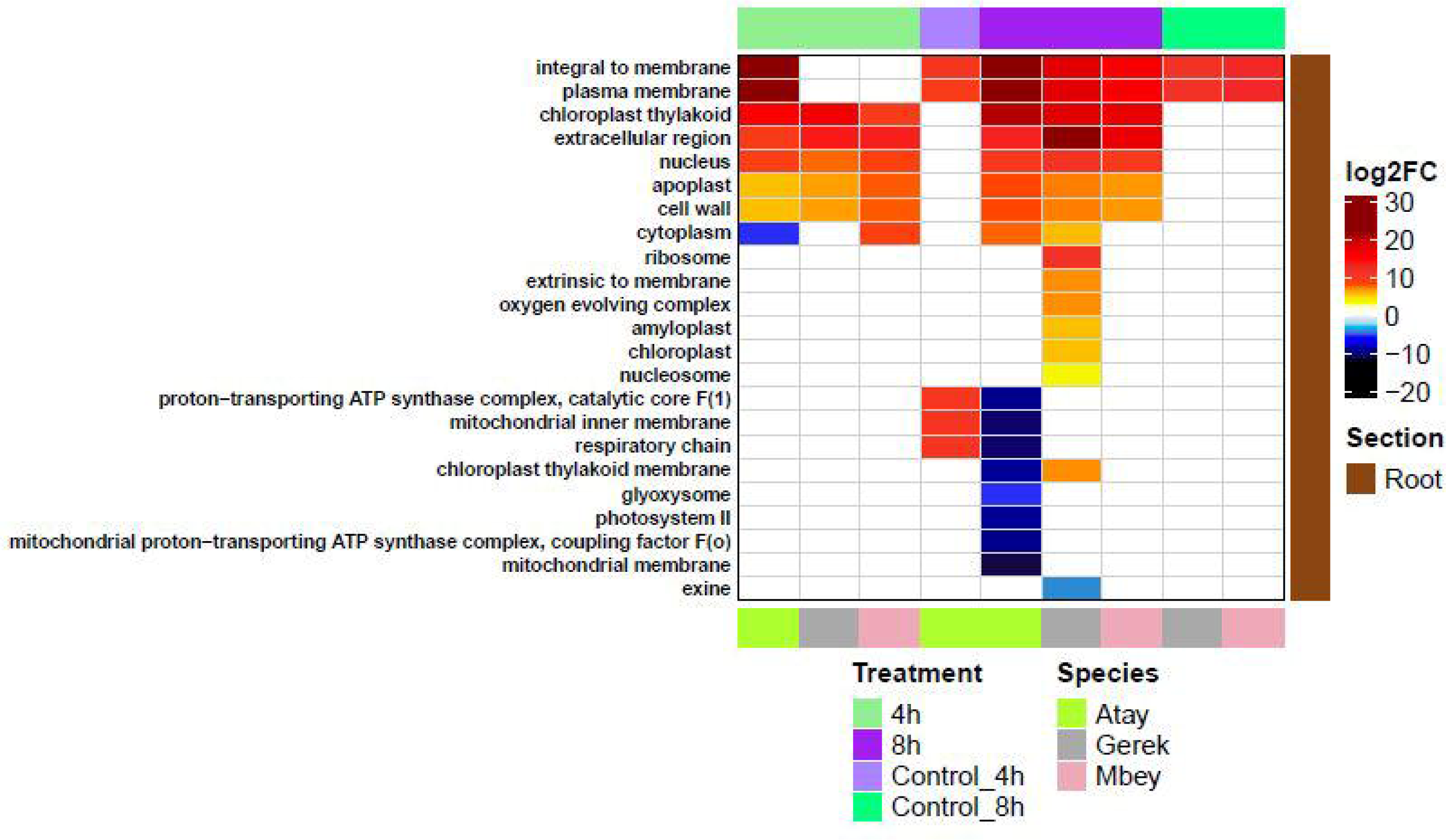
Analysis of cellular component under 4h and 8h drought stress and control groups in leaf and root tissues of the drought-tolerant (Gerek 79, Müfitbey) and sensitive (Atay 85) cultivars. Red and blue colours show the higher and lower expression values, respectively, where Atay 85 is assigned in green, Gerek 79 in grey, and Müfitbey in pink.

**Figure 3.**
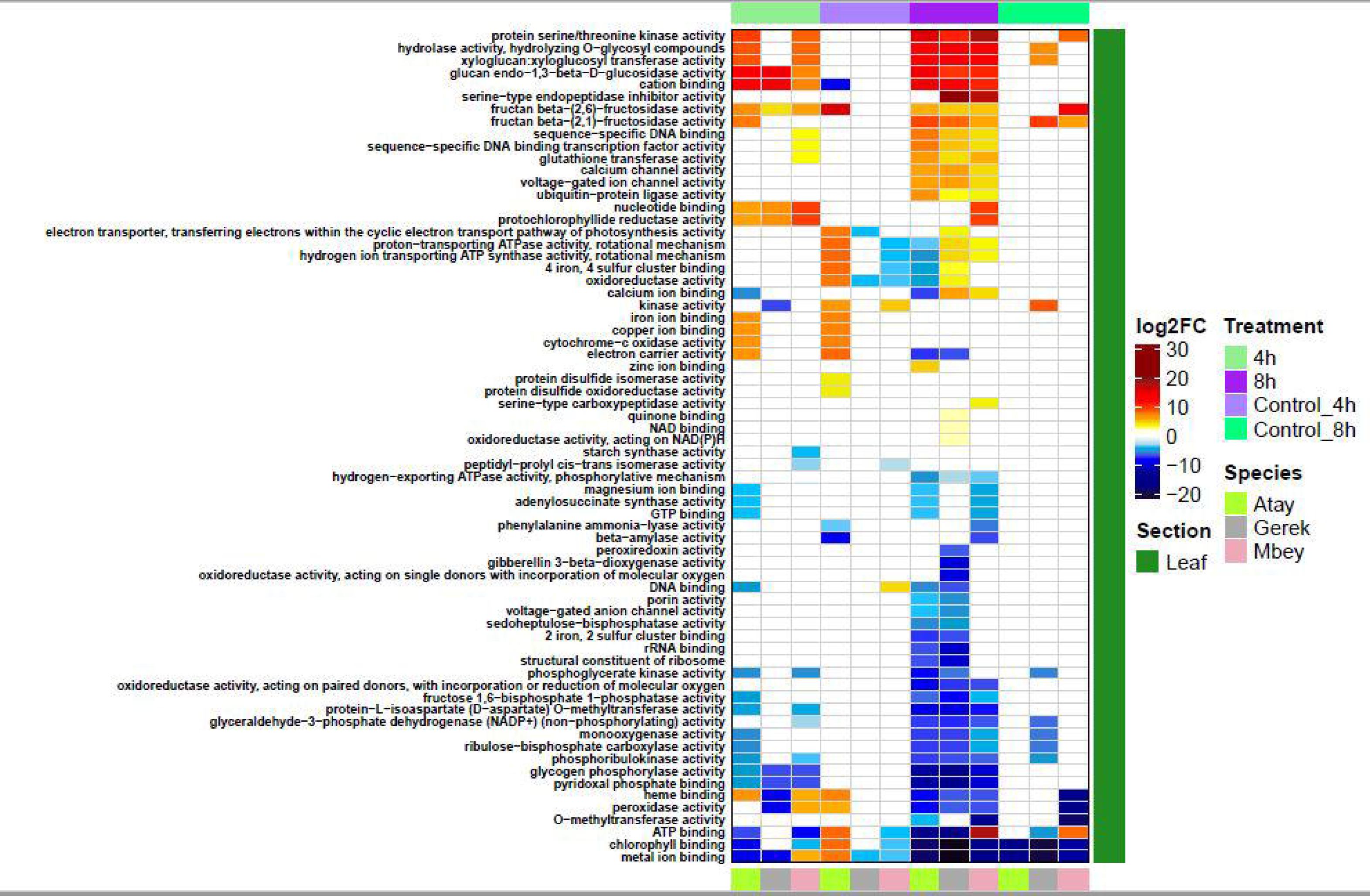
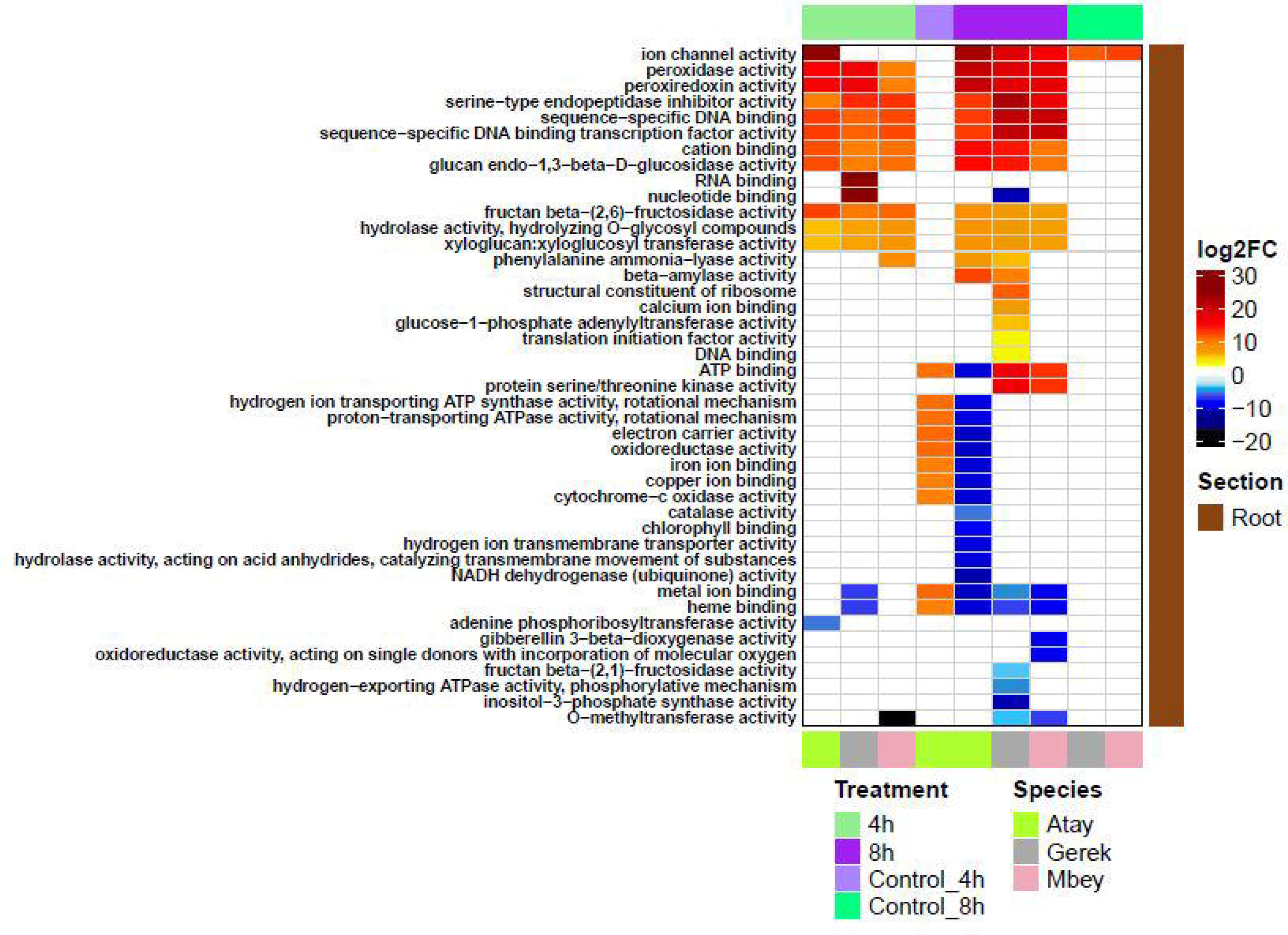
Analysis of Molecular function under 4h and 8h drought stress and control groups in leaf and root tissues of the drought-tolerant (Gerek 79, Müfitbey) and sensitive (Atay 85) cultivars. Red and blue colours show the higher and lower expression values, respectively, where Atay 85 is assigned in green, Gerek in grey, and Müfitbey in pink.

### Validation of DGEs under Drought Stress using qRT-PCR

We have selected eight drought-related genes randomely from the DEG analysis to investigate their expression level using pRT-PCR to confirm our RNA-seq data. Although the fold-changes varied between the RNA-Seq and qRT-PCR analyses, the overall qRT-PCR expression profile of most of the genes agreed with the RNA-Seq profile, indicating the reliability of the RNA-Seq data. To validate the RNA-seq data, differentially expressed genes that might be involved in different stress responses were chosen for further qRT-PCR experiments. The expression of *Probable pectinesterase/ pectinesterase inhibitor 42* (*TaPME42*), *Extensin-like protein* (*TaExLP*), *Germin-like protein 9-1* (*TaGLP9-1*), *Zinc finger CCCH domain-containing protein 36* (*TaZFP36*), *Metacaspase-5* (*TaMC5*), *Phosphoglycerate/bisphosphoglycerate Mutase* (*TaPGM*), *Serine/ threonine protein phosphatase 2A* (*TaPP2CA*), *GIGANTEA* (*TaGI*), *Polyadenylate-binding protein* (*TaRBP45B*), FERRITIN (*TaFER*), *Arogenate dehydratase 5* (*TaADT*), *F-box protein* (*TaFBW2*) genes were investigated in root and leaf tissues of drought-stressed and control plants.

### DEGs Involved in Metal Ion Binding

***Zinc finger CCCH domain-containing protein 36*** *(TaZFP36)* expression was increased in 4h and 8h drought stressed root and leaf tissues of the tolerant cultivar Müfitbey (Figure 4). In contrast, in the drought-sensitive cultivar Atay 85, there was no significant difference between control and drought treated root and leaf tissues (Figure 4).

**Figure 4.**
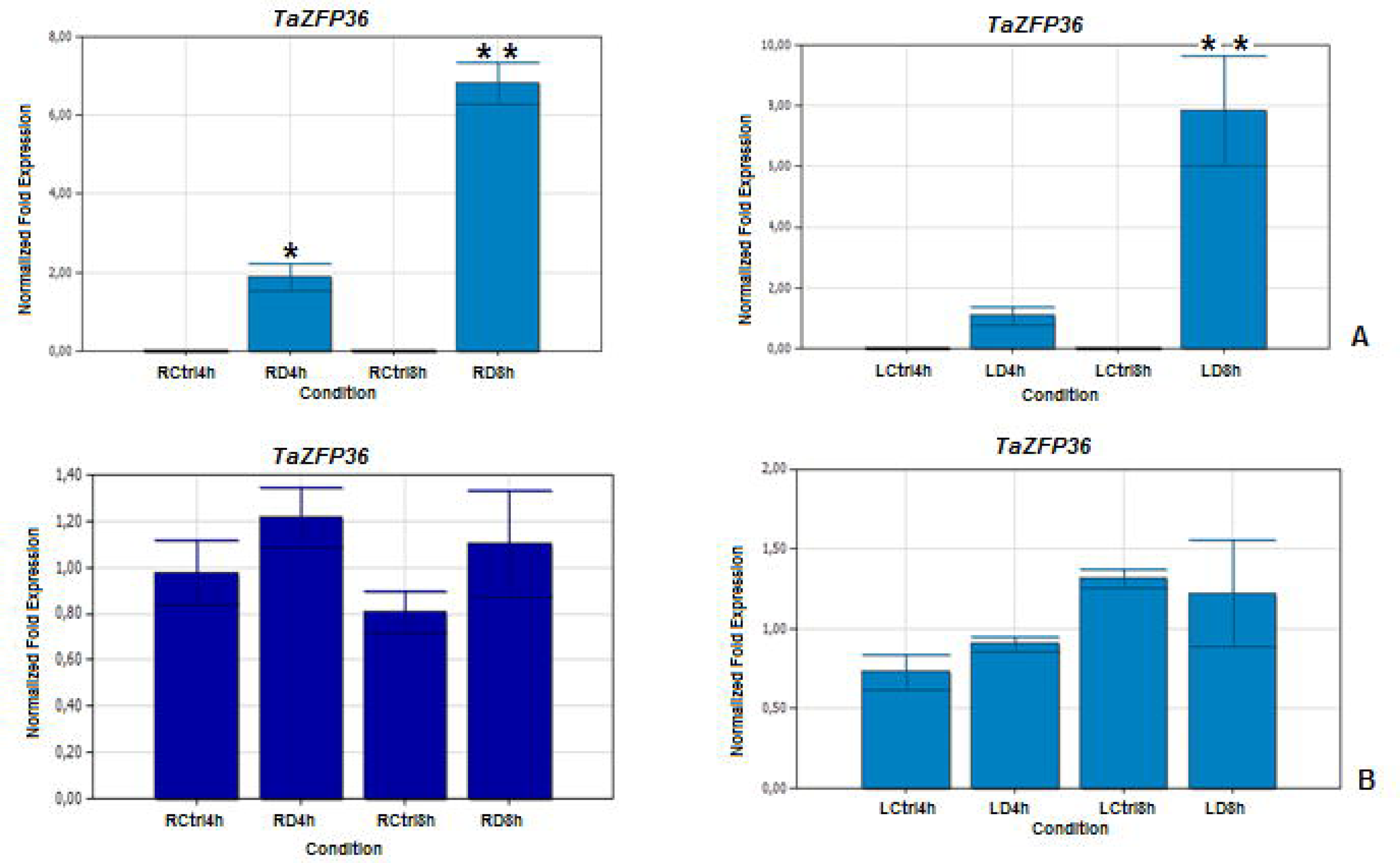
Expression pattern of *Zinc finger CCCH domain-containing protein 36* (*TaZFP36*) gene in 4h and 8h drought-stressed root and leaf tissues. **(A)** Drought-tolerant (Müfitbey) **(B)** Drought**-**sensitive (Atay 85) cultivar. LCtrl, Leaf Control; LD, Leaf Drought; RCtrl, Root Control; RD, Root Drought. Error bars correspond to the standard error of the means.

***Ferritin* (*Fer*)** is involved in ferric iron binding and oxidoreductase activity. In our qRT-PCR experiments, ferritin mRNA expression was found to be differentially expressed in response to drought stress. The expression level of *TaFer* was elevated in 4h and 8h drought stressed leaves in drought-tolerant and drought-sensitive cultivars, especially in 8h drought stressed leaves (Supplementary Figure S10).

### DEGs Involved in Cell Wall Related Genes

***Probable pectinesterase/pectinesterase inhibitor 42 (PME42)***: PME is an enzyme that demethylesterifies a major component of plant cell wall pectins [40]. In our qRT-PCR experiments, an increased level of *TaPME42* was observed in 4h and 8h drought-stressed root and leaf tissue of both the tolerant cultivar Müfitbey, and the drought-sensitive cultivar Atay 85 (Figure 5). In leaf tissue, *TaPME42* expression was also increased in tolerant and drought-sensitive cultivars under different drought stresses (Figure 5).

**Figure 5.**
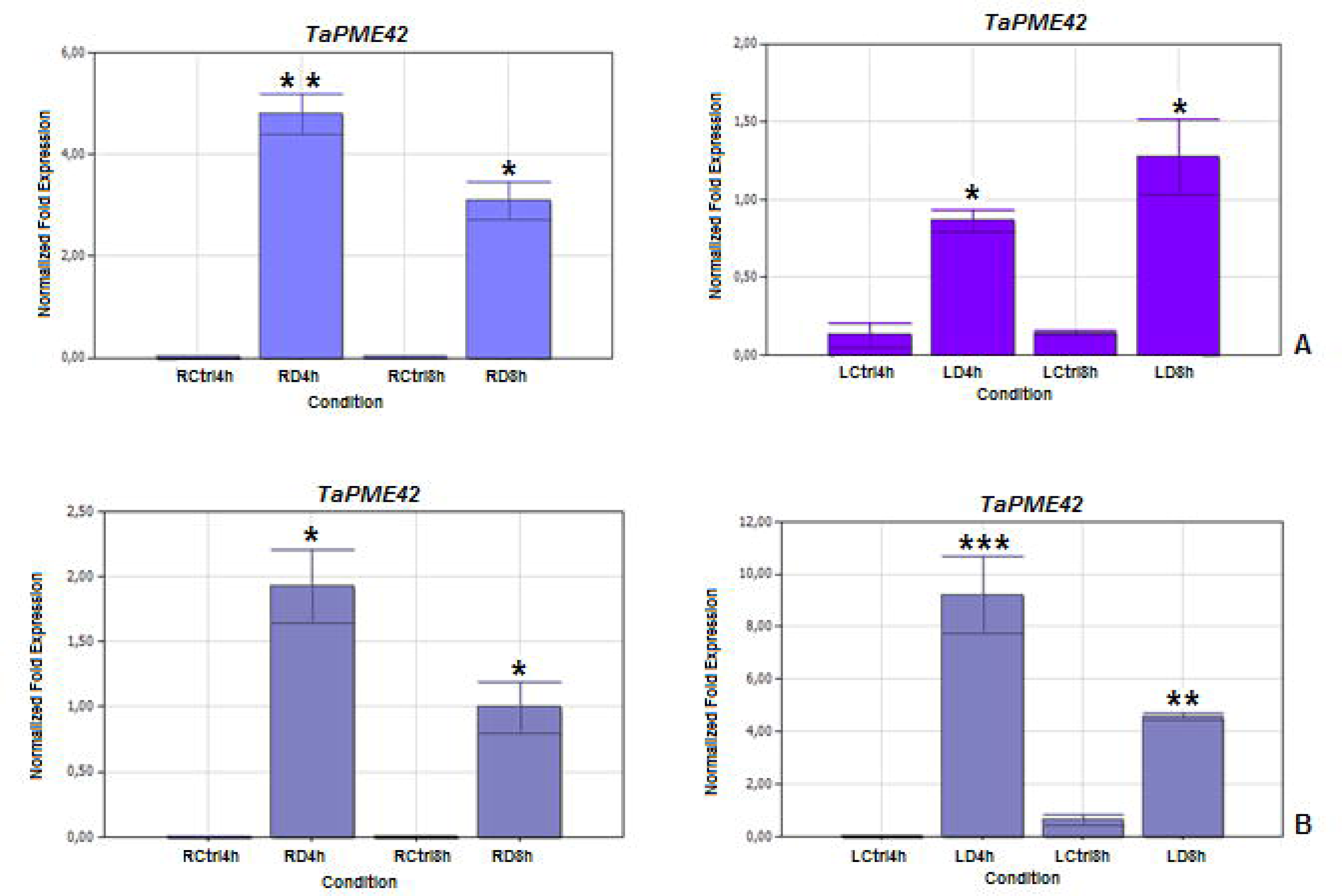
Expression pattern of *pectinesterase/pectinesterase inhibitor 42* (*TaPME42*) gene in 4h and 8h drought-stressed root and leaf tissues. **(A)** Drought-tolerant (Müfitbey), **(B)** Drought-sensitive (Atay 85) cultivars. LCtrl, Leaf Control; LD, Leaf Drought; RCtrl, Root Control; RD, Root Drought. Error bars correspond to the standard error of the means.

***Extensin-like protein (ExLP),*** cell wall extensin is a member of the family of hydroxyproline-rich glycoproteins (HRGPs) which are among the most abundant proteins present in the cells of higher plants [41]. In our qRT-PCR experiments, drought stress caused elevated expression levels of genes coding for extensin-like proteins in roots. Maximum *TaExLP* expression was observed in 4h drought-stressed root tissues of tolerant and drought-sensitive cultivars (Figure 6). In contrast, different expression patterns were observed between the leaf tissues of tolerant and drought-sensitive cultivars. In the drought-sensitive Atay 85 cultivar, the highest TaExLP expression was evident within 4 hours of drought-stressed leaf tissues, with no significant variation observed after 8 hours of stress (as shown in Figure 6). In contrast, the tolerant cultivar Müfitbey exhibited a reduced expression level of this gene in the drought-stressed leaf tissues (as depicted in Figure 6).

**Figure 6.**
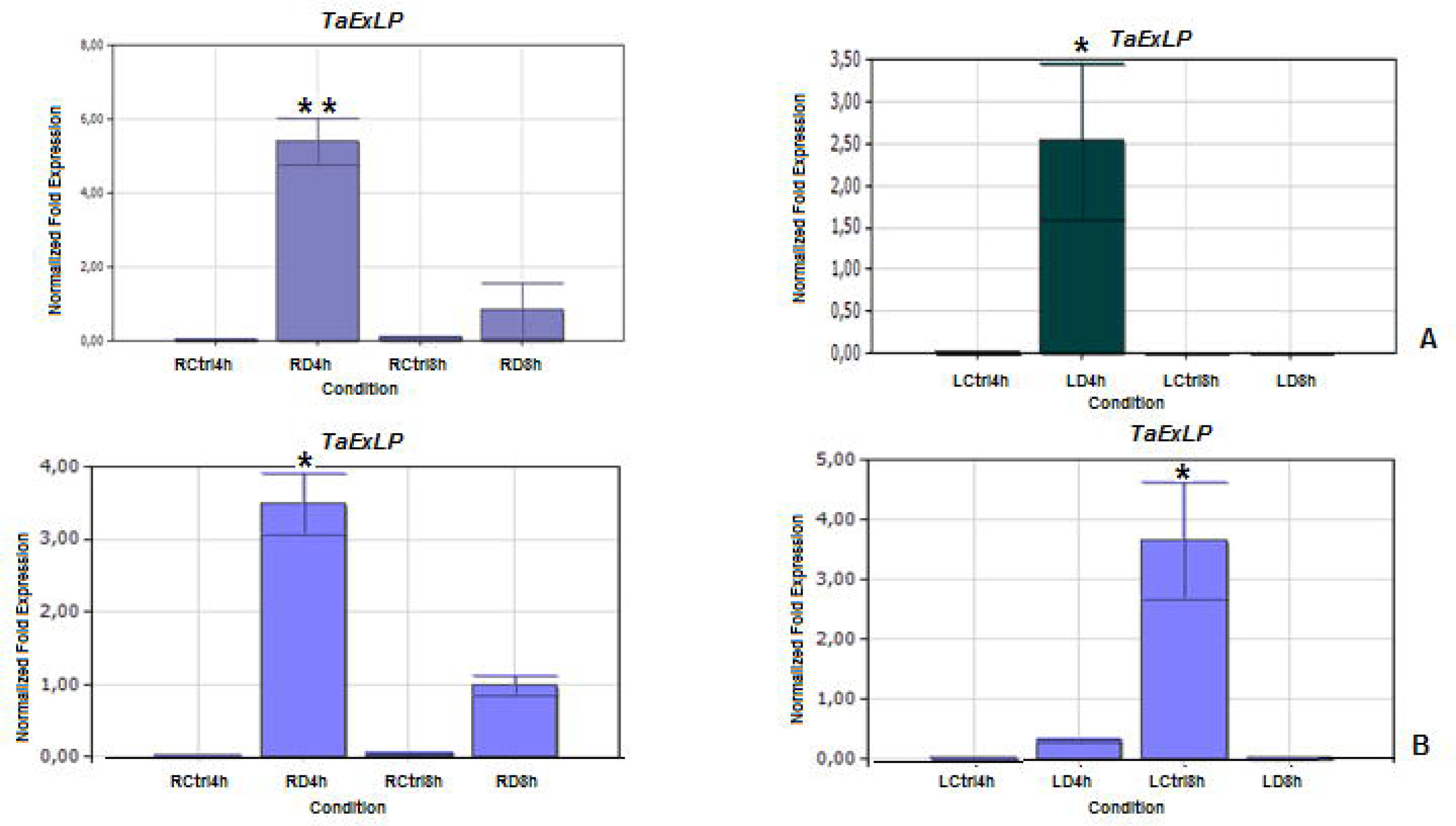
Expression pattern of *Extensin-like protein* (*TaExLP*) gene in 4h and 8h drought-stressed root and leaf tissues. **(A)** Drought-tolerant (Müfitbey), **(B)** Drought-sensitive (Atay 85) cultivars. LCtrl, Leaf Control; LD, Leaf Drought; RCtrl, Root Control; RD, Root Drought. Error bars correspond to the standard error of the means.

***Germin-like protein 9-1 (GLP):*** Germins and GLPs are involved in many processes that are important for plant development and defense mechanisms [42, 43]. We observed that *TaGLP 9-1* was induced in both drought-tolerant and drought-sensitive cultivars in 4h and 8h drought-stressed root tissues (Figure 7). In leaf tissues of the sensitive cultivar, there was no dramatic difference between control and drought-stressed tissues. In contrast, elevated levels of expression were observed in 4h and 8h drought-stressed leaf tissues of the drought-tolerant cultivar Müfitbey (Figure 7).

**Figure 7.**
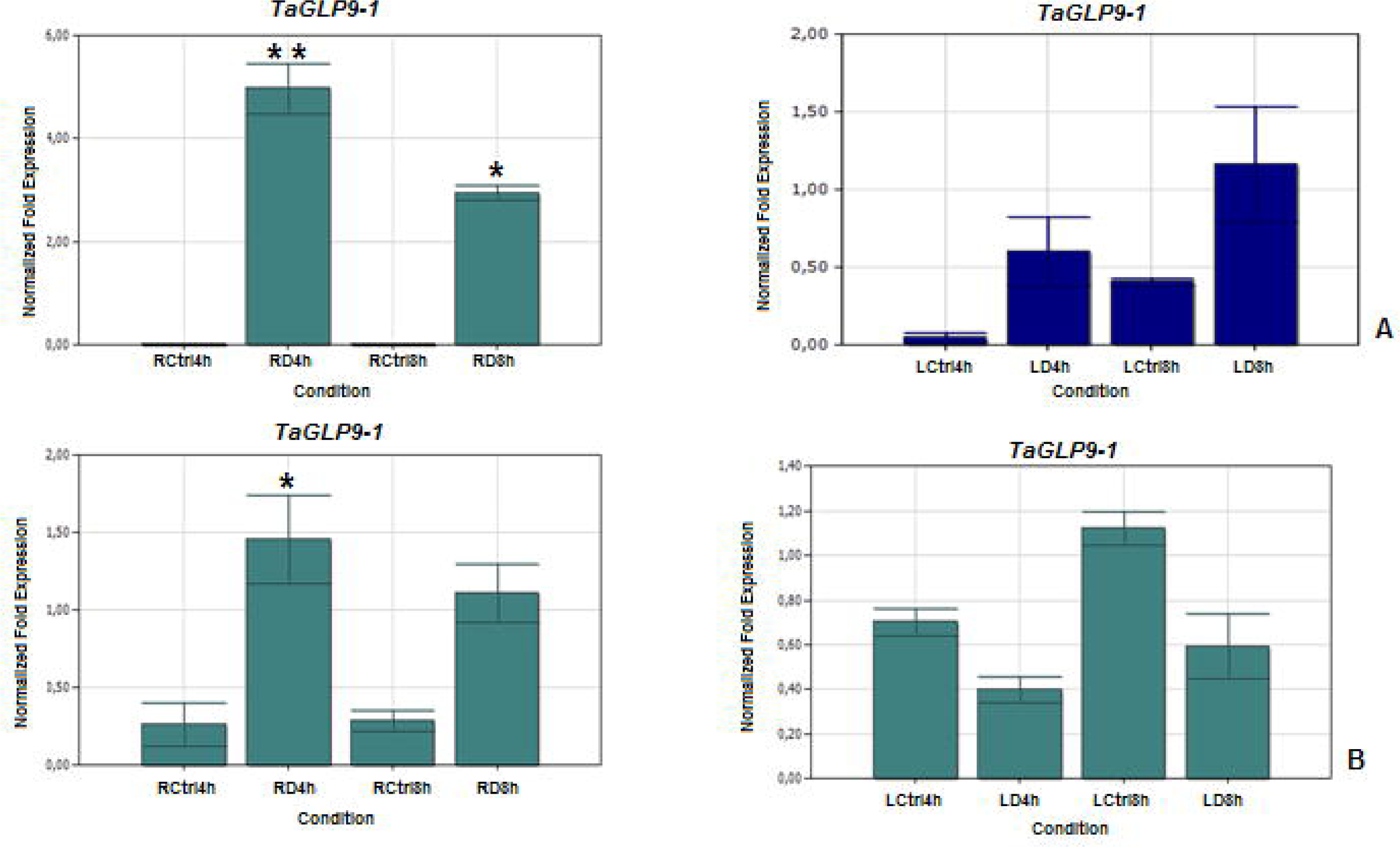
Expression pattern of *Germin-like protein 9-1* (*TaGLP 9-1*) gene in 4h and 8h drought-stressed root and leaf tissues. **(A)** Drought-tolerant (Müfitbey), **(B)** Drought-sensitive (Atay 85) cultivars. LCtrl, Leaf Control; LD, Leaf Drought; RCtrl, Root Control; RD, Root Drought. Error bars correspond to the standard error of the means.

### DEGs Involved in Defense Response Proteins

***Metacaspase -5*** (*MC5*) induces Programmed Cell Death (PCD), an indispensable process in plant and animal immune systems that serves to eliminate cells and/or tissues and recycle nutrients from these tissues to the rest of the organism [44]. RNAseq data showed the expression level of *TaMC5* was elevated in root and leaf tissues of tolerant cultivar Müfitbey after 8h drought stress. However, in qRT-PCR experiments, the *TaMC5* expression level in tolerant cultivars was increased in 4h and 8h of drought stress; leaf tissues showed no significant increase (Figure 8). In qRT-PCR analysis of the sensitive cultivar Atay 85, the expression level of *TaMC5* was not significantly affected by droght stress in either root or leaf root tissues (Figure 8).

**Figure 8.**
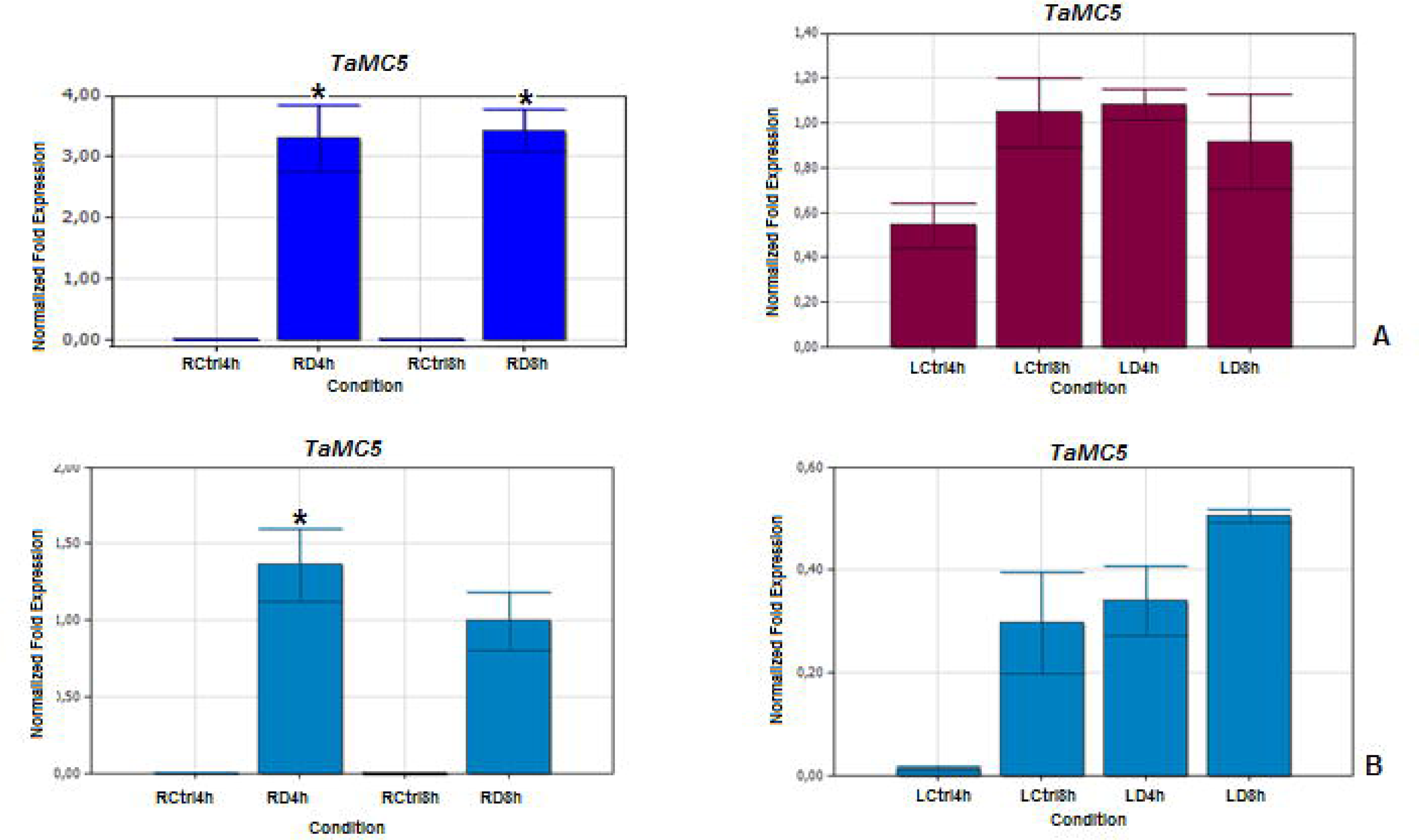
Expression pattern of *Metacaspase-5* (*TaMC5*) in 4h and 8h drought-stressed root and leaf tissues. **(A)** Drought-tolerant (Müfitbey), **(B)** Drought-sensitive (Atay 85) cultivars. LCtrl, Leaf Control; LD, Leaf Drought; RCtrl, Root Control; RD, Root Drought. Error bars correspond to the standard error of the means.

***Arogenate Dehydratase 5 (ADT-5)*** expression was increased in leaf tissues after 4 h drought stress in both sensitive and tolerant cultivars (Supplementary Figure S11). In addition, after 8 h of drought stress, the expression level of *TaADT-5* in leaf tissue of the drought-tolerant cultivar Müfitbey was very significantly increased by eight-fold. Conversely, in the sensitive cultivar, there was a two-fold decreased level in expression level of *ADT-5* in 8 h drought-stressed leaf tissues (Supplementary Figure S11).

### DEGs Involved in Carbohydrate degradation

***Phosphoglycerate/bisphosphoglycerate mutase (PGM)*** catalyzes reactions involving the transfer of groups between the three carbon atoms of phosphoglycerate [45]. *TaPGM* expression was increased in root tissues of the sensitive cultivar after 4 and 8 hours of drought stress. It’s expression level in the tolerant cultivar was significantly elevated in 4h drought-stressed leaf tissue, while difference in the expression level was not observed in 8h drought stressed leaves (Figure 9).

**Figure 9.**
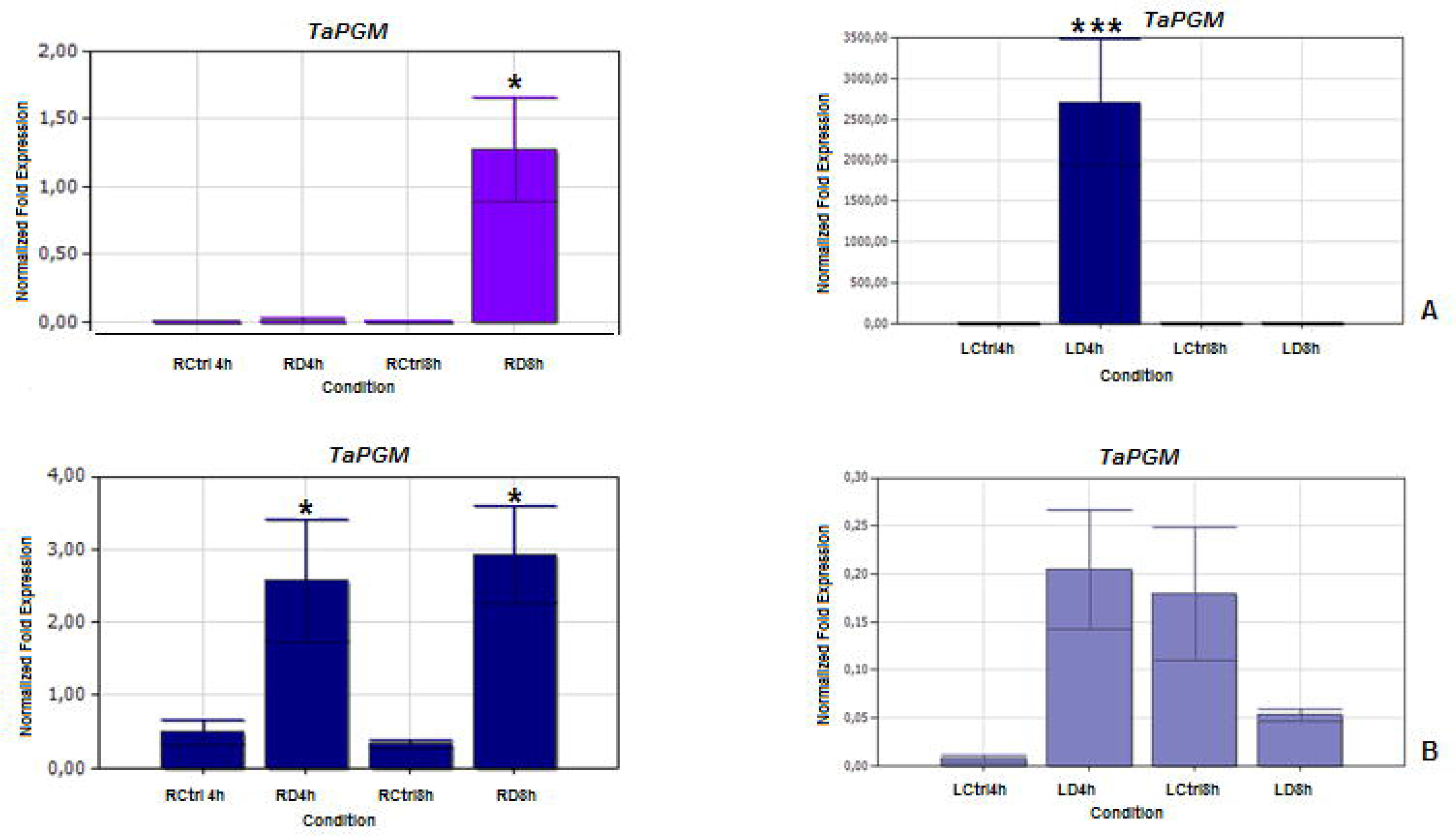
Expression pattern of *Phosphoglycerate/bisphosphoglycerate mutase* (*TaPGM*) in 4h and 8h drought-stressed root and leaf tissues. **(A)** Drought-tolerant (Müfitbey), **(B)** Drought-sensitive (Atay 85) cultivars. LCtrl, Leaf Control; LD, Leaf Drought; RCtrl, Root Control; RD, Root Drought. Error bars correspond to the standard error of the means.

### DEGs Involved in ABA-related gene expression

***Serine/threonine protein phosphatase 2A (PP2A)*** regulates beta-oxidation of fatty acids and protoauxins in peroxisomes by dephosphorylating peroxisomal beta-oxidation-related proteins [46]. *TaPP2CA* expression was significantly increased in both leaf and root tissues of tolerant and sensitive cultivars after 4 and 8 hours drought stress. In root and leaf tissues of the tolerant cultivar and leaf tissues of the sensitive cultivar, maximum expression was observed in 4h drought-stress, with lower, though still significant, expression levels after 8h drought stress (Figure 10). However, in the root tissues of the sensitive cultivar, expression levels were similarly increased after both 4h and 8h drought stress (Figure 10).

**Figure 10.**
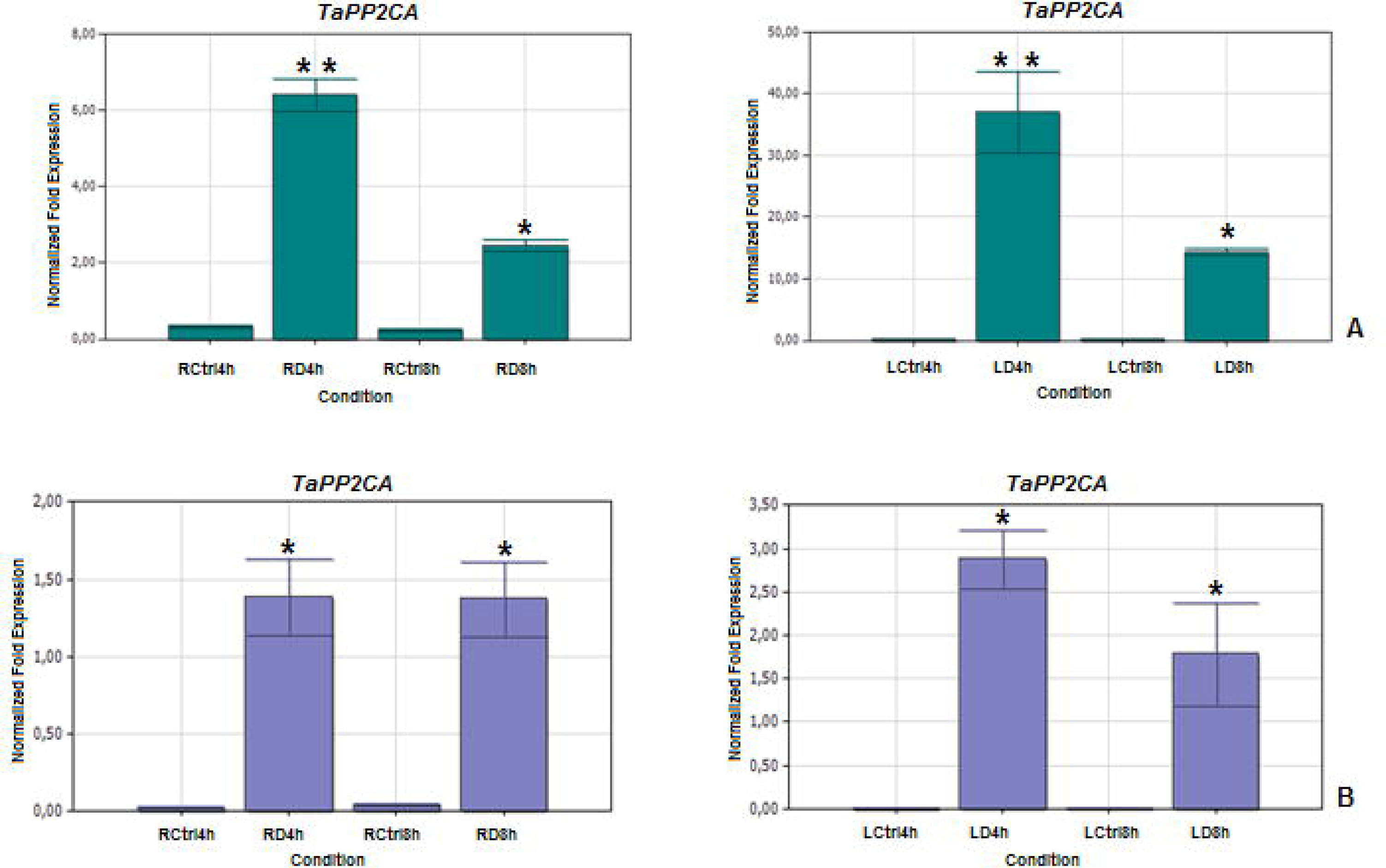
Expression pattern of *Serine/threonine protein phosphatase 2A (TaPP2CA)* in 4h and 8h drought-stressed root and leaf tissues. **(A)** Drought-tolerant (Müfitbey), **(B)** Drought-sensitive (Atay 85) cultivars. LCtrl, Leaf Control; LD, Leaf Drought; RCtrl, Root Control; RD, Root Drought. Error bars correspond to the standard error of the means.

### DEGs Involved in Regulation of Photoperiodism and Flowering

***Protein GIGANTEA (GI)*** is involved in the regulation of circadian rhythm, photoperiodic, and phytochrome B signaling and flowering [47]. In leaf tissues of the drought-tolerant cultivar Müfitbey, expression of *TaGI* was reduced after 8h drought stress, but conversely, in leaf tissues of the sensitive cultivar, expression was increased after 4 h of drought stress (Figure 11). Significant changes in expression in response to drought were not observed in the root tissues of either the tolerant or the sensitive cultivars (Figure 11).

**Figure 11.** Expression pattern of *GIGANTEA* (*TaGI*) in 4h and 8h drought-stressed root and leaf tissues. **(A)** Drought-tolerant (Müfitbey), **(B)** Drought-sensitive (Atay 85) cultivars. LCtrl, Leaf Control; LD, Leaf Drought; RCtrl, Root Control; RD, Root Drought. Error bars correspond to the standard error of the means.

***Polyadenylate-binding protein (RBP45B):*** Heterogeneous nuclear ribonucleoprotein (hnRNP) binds the poly(A) tail of mRNA and is probably involved in some steps of pre-mRNA maturation. Expression of *TaRBP45B* was found to be induced by 4h and 8h drought-stress in root tissues of both tolerant and sensitive cultivars (Figure 12). On the other hand, in leaf tissues, a significant increase of expression level was observed after 4h of drought stress in both the tolerant cultivar Müfitbey and the sensitive cultivar Atay 85, but neither cultivar showed a significant change in expression level from the control after 8 h of drought stress (Figure 12).

**Figure 12.**
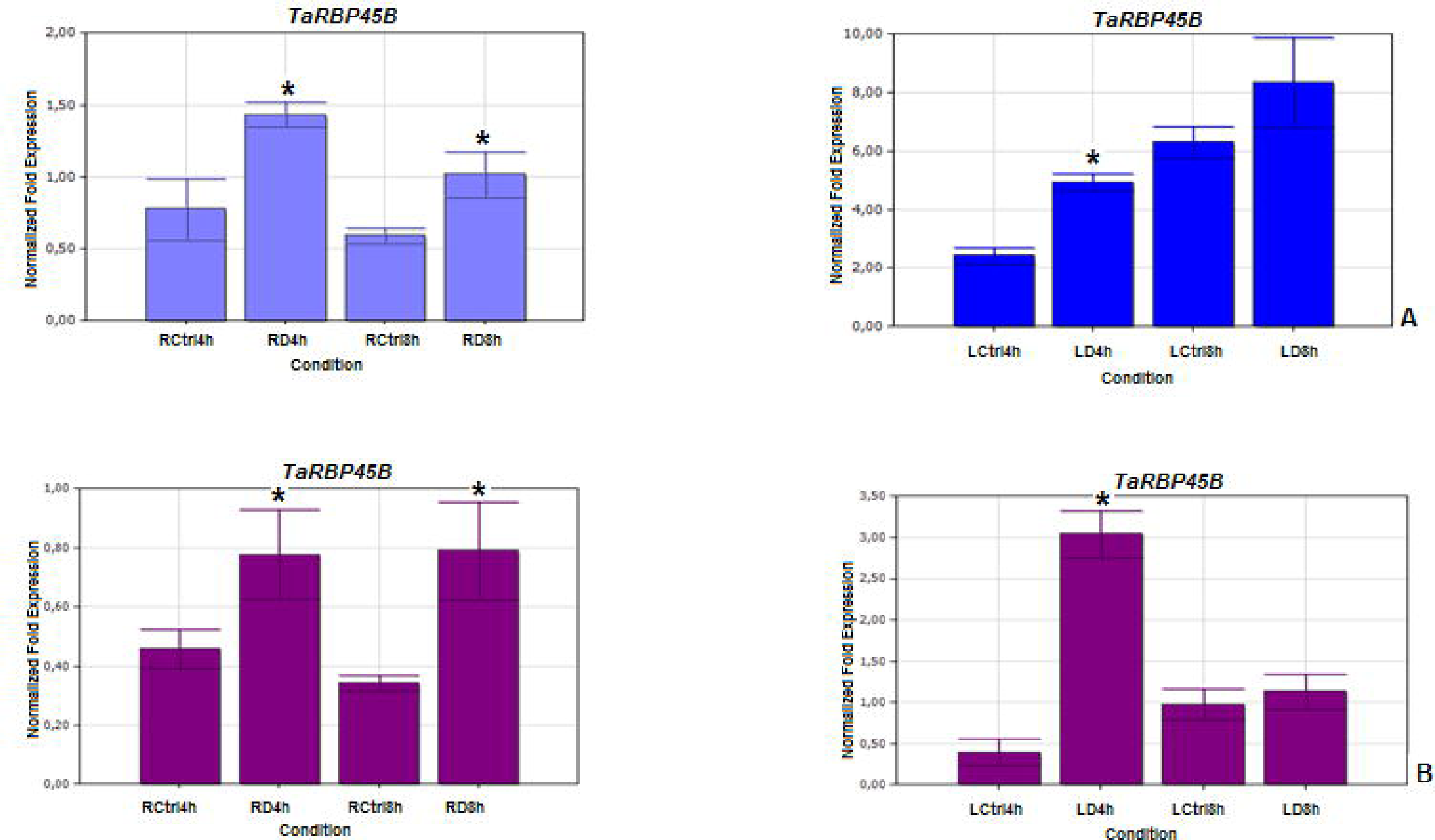
Expression pattern of *Polyadenylate-binding protein (TaRBP45B)* in 4h and 8h drought-stressed root and leaf tissues. **(A)** Drought-tolerant (Müfitbey), **(B)** Drought-sensitive (Atay 85) cultivars. LCtrl, Leaf Control; LD, Leaf Drought; RCtrl, Root Control; RD, Root Drought. Error bars correspond to the standard error of the means.

## DISCUSSION

Drought stress has a severe impact on plant growth and can lead to significant reductions in wheat yields, particularly in cultivated areas. To comprehensively understand the drought stress mechanism in hexaploid wheat, it is crucial to study gene expression in both tolerant and sensitive genotypes. While there have been various studies on drought stress-related transcriptome analysis in different crop plants [48], the specific mechanisms in tolerant and sensitive *T. aestivum* cultivars have not been extensively investigated.

In this study, we aimed to provide a comprehensive understanding of drought stress-related gene expression in response to drought stress in two different drought-tolerant and one drought-sensitive *T. aestivum* cultivars. Our findings revealed distinct physiological and molecular responses in root and leaf tissues under drought stress, with variations observed at both 4-hour and 8-hour time points. These responses also differed from their respective control groups.

In leaf tissue, a noticeable trend of decreased gene expression was observed for cellular processes such as protein refolding and cellular metabolic processes like photorespiration as drought stress duration increased (8 hours) in all three cultivars. The comparison of transcriptome profiling across all cultivars provided valuable insights into the complexity of the drought stress response at the molecular level. Our RNA-seq data indicated that metabolic processes related to gene expression were predominantly activated in response to 4-hour and 8-hour drought stress.

Our results further highlighted that drought-tolerant cultivars (Müfitbey and Gerek 79) exhibited increased expression levels of genes related to protein binding, metabolic processes, and cellular functions, indicating their ability to adapt better to drought stress compared to the sensitive cultivar Atabey 85. Similar studies on *Cucumis sativus* L. plants exposed to drought stress also reported significant increases in gene expression, especially in metabolic processes, membrane-related functions, and catalytic activity [49].

Transcription factors (TFs) have been considered putative candidate genes capable of regulating gene expression in response to different stresses [50]. By binding directly to the promoters of target genes in a sequence-specific manner, they activate or suppress the activation of downstream genes [51]. For that reason, the identification and evaluation of TF genes related to stress tolerance are essential for molecular improvement in different breeding programs. In the sensitive cultivar, we detected more than 25 differentially expressed TFs in leaf tissues under 4- hour and 8-hour drought stress, while only four TFs were identified in root tissues. In contrast, the tolerant cultivar exhibited more than 80 different TF transcripts in both leaves and roots after 4 hours of drought stress, with this number decreasing to 18 after 8 hours of drought stress. These findings underscore the role of TFs in drought tolerance and suggest that multiple TFs contribute to the mechanism of drought resistance.

The expression level of genes related to hydrogen peroxide catabolic processes, photorespiration, glycolysis, and photosystem II stabilization decreased in leaf tissues under 8-hour drought stress, while genes associated with carbohydrate metabolic processes, defense responses, and cellular glucan metabolic processes increased during both 4-hour and 8-hour drought stress in leaf and root tissues. In the sensitive cultivar, the expression levels of genes involved in oxidative phosphorylation, aerobic respiration, ATP hydrolysis and synthesis, and electron transport chain were decreased by 8-hour drought stress in root tissue (Figure 1).

### Metal Ion Binding Plays a Role in Drought Response

Our study revealed significant gene expression related to metal ion binding, heme binding, 2 iron 2 sulfur cluster binding, zinc ion binding, iron ion binding, and copper ion binding proteins in both leaf and root tissues under drought stress in wheat. These metal-ion binding proteins, such as AtTZF2, AtTZF3, and AtTZF1, have well-conserved roles in controlling plant growth, development, and stress responses [52]. A genome-wide analysis of CCCH zinc finger proteins (TZFs) in *Arabidopsis* has revealed 11 members that contain a plant-specific TZF motif [52, 53]. *AtTZF1- AtTZF*6 and *AtTZF9* are involved in ABA response, seed germination, and Pathogen-Associated Molecular Pattern (PAMP)-triggered immune response [54]. Most TZFs can localize to processing bodies (PBs) and stress granules (SGs) and play important roles in post-transcriptional regulation and epigenetic modulation of gene expression [54]. Reverse genetic analyses indicate that *AtTZF1* acts as a positive regulator of ABA response, and a negative regulator of GA response, in part by differential regulation of ABA and GA responsive genes. *AtTZF1* gain-of-function plants are superior to wild type (WT) plants in cold and drought tolerance [55]. *AtTZF2* and *AtTZF3*, two close homologs of *AtTZF1*, appear to play similar roles in controlling plant growth, development, and stress responses [56].

Our results suggest that *TaZFP36* is important for drought tolerance. *TaZFP36* expression was increased in 4h and 8h drought stressed root and leaf tissues of tolerant cultivars. On the other hand, in the sensitive cultivar Atay 85, there was no significant difference between control and drought treated root and leaf tissues. The fact that this gene shows high expression level in drought-resistant plants but does not show any expression difference in sensitive plants suggests that *TaZFP36* may have an important role in drought tolerance mechanism. Our results seem to be compatible with *AtTZF1* gain-of-function studies performed in *Arabidopsis*.

Ferritin gene expression was found to be regulated by oxidative stress, affecting both gene expression and Iron Regulatory Protein activity [57]. Different abiotic stresses, such as ozone or ethylene treatment, iron overload, or impaired photosynthesis, induce ferritin accumulation in chloroplasts [58, 59, 60]. Our qRT-PCR experiments demonstrated differential expression of *TaFer* in response to drought stress in leaf tissues of both tolerant and sensitive cultivars, highlighting the role of oxidoreductase activity in drought stress responses.

### Cell Wall Proteins Clearly Play a Role in Drought Response

Different cell wall protein related genes such as *Beta-galactosidase 1*, *Glucose-6- phosphate/phosphate-translocator*, *Leucine-rich repeat extensin-like protein 4*, *Leucine-rich repeat extensin-like protein 6*, *Germin Like Protein 9-1*, lignin biosynthesis related genes were identified from DEG data. We selected *PME inhibitor* 49 (*TaPME49*), *Extensin-like protein (TaExLP),* and *Germin Like Protein 9-1 (TaGLP9-1)* genes because of their high expression level by drought stress in wheat. The expression level of *PME inhibitor 49* and *Extensin-like protein* genes were increased by drought stress. PME is a demethylesterification of cell wall pectins [40] and has been reported to play a role in different developmental processes, such as hypocotyl elongation [61] and cell differentiation [62]. *TaPME* expression level was elevated in 4h and 8h drought-stressed leaf tissues of tolerant and sensitive cultivars, respectively. In contrast, decreased expression level was observed in 4h and 8h drought-stressed root tissue of tolerant and sensitive cultivars.

*ExLP* is a member of the family of hydroxyproline-rich glycoproteins (HRGPs), which are the most abundant proteins, present in the cell wall of higher plants [41]. Drought stress triggers the expression level of *TaExLP* in roots. Maximum expression of this gene was observed in 4h drought-stressed root tissues of tolerant and sensitive cultivars. The increased level of expression of this gene in the sensitive cultivar in the early period of drought and the suppression in the tolerant cultivar suggest that this gene might be one of the drought susceptibility genes.

Germins and GLPs are involved in many processes that are important for plant development and defense mechanisms [42]. Involvement of significant number of GLPs has been shown in abiotic stress conditions such as salt stress [63], aluminum stress [64] and drought stress [65, 66]. Overexpression was also observed when attacked by fungal pathogens, bacteria, and viruses [67, 68, 69]. GLPs influence plant defense systems because of their generation of reactive oxygen species. They are targeted at the cell wall and apoplast, and some members related to the barley *HvGER4* subfamily exhibit superoxide dismutase activity [70]. The increase expression level of in tolerant wheat cultivar under drought stress suggests that *TaGLP 9-1* is related to drought tolerance in bread wheat.

### Defense Response Proteins in Drought Stress

Defense response related gene expression was increased by 4h and 8h drought-stressed leaf tissues. *Arogenate dehydratase 5* (*ADT5*) plays an important role in lignin biosynthesis [71]. In *Arabidopsis* genome, there are six *ADT* genes designated as *ADT1*–*ADT6* and are ubiquitously expressed in various tissues or organs [72]. It has been reported that *ADT1* and *ADT3* play more important roles in sucrose and cold-induced anthocyanin synthesis [73]. Our results show *ADT-5* mRNA level is increased in tolerant cultivars, indicating that this gene may be involved in the drought stress response.

Metacaspases, a family of cysteine proteases induce programmed cell death (PCD) during plant development and defense responses [74]. A total of nine metacaspases has been identified in *Arabidopsis.* In the Genevestigator analysis, gene expressions of *Arabidopsis*, rice, and tomato, metacaspase family in the developmental stages were investigated. mRNA levels of *OsMC2*, *OsMC6*, and *OsMC7* were all induced by temperature stress [75]. In our research, we noted an elevation in the *Metacaspase-5* (*TaMC5*) mRNA levels within the 8-hour drought-stressed root and leaf tissues of the tolerant cultivar Müfitbey. This observation strongly indicates the significance of the *TaMC5* gene in conferring drought tolerance to *T. aestivum*.

**Drought Stress Activates Carbohydrate Degradation-related Genes** Phosphoglycerate/Bisphosphoglycerate Mutase (PGM) facilitates reactions involving the transfer of phosphate groups within the three carbon atoms of phosphoglycerate. It dephosphorylates and activates Actin-Depolymerizing Factor 1 (ADF1), a protein that governs the re-modelling of the actin cytoskeleton [76]. Notably, the expression level of *TaPGM* exhibited a substantial increase after 8 hours of drought stress in the roots of the drought-sensitive cultivar Atay 85 (Figure 9). In contrast, *TaPGM* expression showed a significant increase in 4 hour drought-stressed leaf tissue of the tolerant cultivar, with no differential expression observed in 8-hour drought-stressed leaves (Figure 9). The upregulation of this gene in the sensitive cultivar suggests that this gene may be required for susceptibility to drought stress.

### Involvement of ABA-related Genes in Drought Stress

*Serine/threonine Protein Phosphatase 2A (PP2A)* acts as a negative regulator of ABA signalling and is involved in the regulation of ABA-dependent gene expression and the light-dependent activation of nitrate reductase [77, 78]. In rice (*Oryza sativa*), all catalytic subunit genes (*OsPP2A-1-5*) are upregulated in response to high salinity in leaves [79]. In the same way, salt stress increases mRNA levels of potato *StPP2Ac1, StPP2Ac2a, StPP2Ac2b,* and *StPP2Ac3* in leaves [80]. Furthermore, okadaic acid inhibits the salt stress response in potatoes, indicating a positive regulation by Ser/Thr phosphatases [80]. *TaPP2Ac-1* catalytic subunit transcripts accumulate in seedlings in response to water deficits [80]. Transgenic tobacco plants overexpressing *TaPP2Ac-1* exhibit enhanced drought tolerance, indicating that this PP2A catalytic subunit acts as a positive regulator of salt stress adaptive responses [81].

The maximum *TaPP2CA* expression was detected in drought-stressed leaf and root tissues of tolerant and sensitive cultivar after 4 and 8 hour (Figure 10) indicating its importance in the early stage of the drought tolerance mechanism.

### Regulation of Photoperiodism in Drought Stress

Decreased photosynthesis, light harvesting process, photosystem I stabilization, and photorespiration related gene expression were decreased in 8h drought-stress tolerant and sensitive plants. Protein GIGANTEA is involved in the regulation of circadian rhythm, photoperiodic, phytochrome B signaling, and flowering [82]. It was also reported that *GI* regulation was affected by cold, hydrogen peroxide, blue light, and Karrikin [82, 83]. It stabilizes *Adagio protein 3* (*ADO3*) and the circadian photoreceptor ADO1/ZTL and regulates ‘CONSTANS’ (CO) in the long-day flowering pathway. It is known that GI provides high salinity tolerance through interaction with the protein kinase SALT OVERLY SENSITIVE 2 (SOS2) and induces EARLY FLOWERING (ELF) under drought stress conditions [84, 85]. Mutations in GI increase resistance to oxidative stress and freezing through upregulation of CDF expression [86, 87]. The biochemical mechanism of GI in the stress response have not been elucidated in detail. In our qRT-PCR, *TaGI* expression was decreased in 8h drought-stressed leaf tissues of the tolerant cultivar Müfitbey (Figure 11) demonstrating the negative effect of this gene on the drought tolerance mechanism, which is compatible with the *GI* gene knockout studies.

Polyadenylate-binding protein RBP45B (RNA-binding protein 45) is related to heterogeneous nuclear ribonucleoprotein (hnRNP)-protein binding the poly (A) tail of mRNA and is likely to be involved in some steps of pre-mRNA maturation, and translation initiation during stress conditions in plants. The upregulation of RBPs in response to plant adaptation to abiotic stress (salt, drought, heat, cold, ozone, hypoxia and flooding) implying its importance for abiotic stress tolerance [88]. *TaRBP45B* was found to be induced by 4h and 8h drought-stressed root tissues. On the other hand, in leaf tissue, significant differences were obtained in 4h stressed leaves of tolerant and sensitive cultivars (Figure 14). Increased level of expression of *TaRBP45B* indicates its positive role in drought tolerance mechanism, in line with the other abiotic stress studies of RBPs.

Understanding abiotic stress tolerance is an indispensable way of adapting to environmental conditions. This study contributes to the identification and illumination of the complex drought stress mechanism. Functional characterisation of genes that play a role in the complex drought-response in wheat will be helpful for developing wheat varieties that are more productive with less water.

## DATA AVAILABILITY STATEMENT

The datasets presented in this study can be found in online repositories. The names of the repository/repositories and accession number(s) can be found in the article.

## AUTHOR CONTRIBUTIONS

Conceptualization, B.C.K.; methodology, B.C.K, I.T., and A.H.S.Ç.; data analysis, O.U.S., R.F., and B.Ö.; validation, A.H.S.Ç., Y.Y, S.O.; formal analysis, M.T.; data curation, M.T.; writing-original draft preparation, B.C.K.; writing-review and editing, M.T.; supervision, B.C.K. All authors have read and approved the submitted version of the manuscript.

## ACKNOWLEDGEMENT

We would like to acknowledge Agriculture and Forestry Translational Zone Agricultural Research Institute (Eskişehir-Turkey) and the Ankara Agricultural Research Institute (Turkey) for providing bread wheat cultivars used in this study. We also thank to Dr. Alison Woods-Tör for critically reading the manuscript. We would also like to thank Fatih Karakaya and Dr. H. Aslı Yalçın-Ercan for their technical support.

## Supplementary Figure Legends

**Supplementary Figure S1.** Four-week old bread wheat cultivars grown in plant growth room for initial screening. **A)** untreated (control), **B)** drought stress induced plants.

**Supplementary Figure S2.** Relative water content (RWC) measurements of various wheat cultivars after 10d drought stress.

**Supplementary Figure S3.** Shock dehydration drought stress induction in Gerek 79, Atay 85 and Müfitbey cultivars. **(A)** 4 h and **B)** 8 h after removal from the hydroponic culture.

**Supplementary Figure S4.** Gene Ontology (GO) Biological Process of Root Tissues

**Supplementary Figure S5.** Gene Ontology (GO) Biological Process of Leaf Supplementary

**Supplementary Figure S6.** Gene Ontology (GO) Cellular Component of Root Tissue

**Supplementary Figure S7.** Gene Ontology (GO) Cellular Component of Leaf Tissue

**Supplementary Figure S8.** Gene Ontology (GO) Molecular Function of Root Tissue

**Supplementary Figure S9.** Gene Ontology (GO) Molecular Function of Leaf Tissue

**Supplementary Figure S10.** The Expression pattern of ***Ta Ferritin*** in 4h and 8h drought stressed leaf tissues of drought-tolerant (Müfitbey) and sensitive (Atay 85) cultivars. LCtrl, Leaf Control; LD, Leaf Drought. Error bars correspond to the standard error of the means. A. Müfitbey, B. Atay 85.

**Supplementary Figure S11.** The expression pattern of ***Arogenate dehydratase 5* (*TaADT*)** in drought stressed and control leaf tissues of drought-tolerant (Müfitbey) and sensitive (Atay 85) cultivars. LCtrl, Leaf Control; LD, Leaf Drought. Error bars correspond to the standard error of the means. A. Müfitbey, B. Atay 85.

## Supplementary Tables

**Supplementary Table 1.**
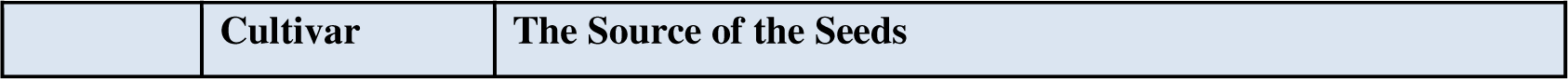
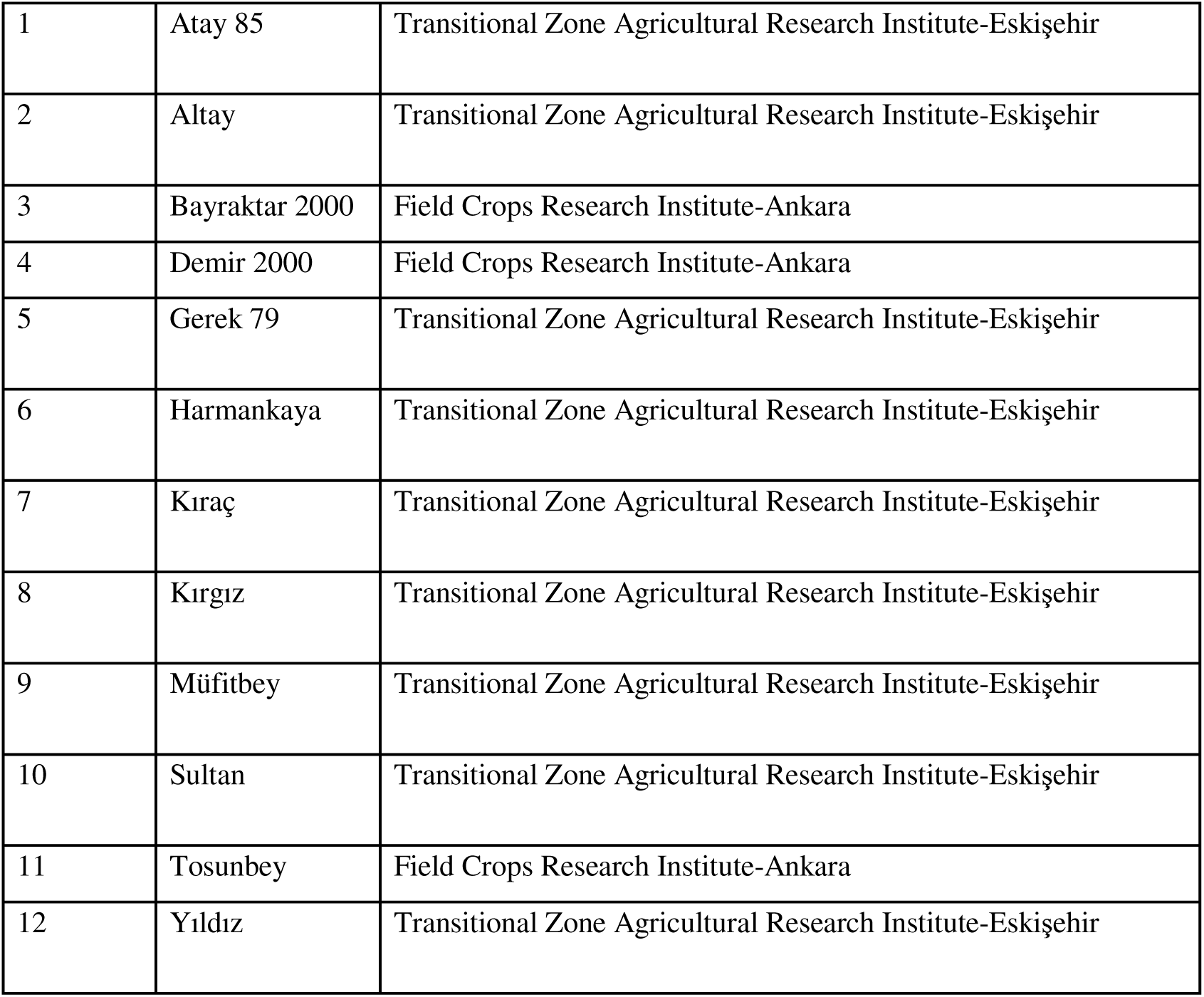
List of cultivars used.

**Supplementary Table 2.**
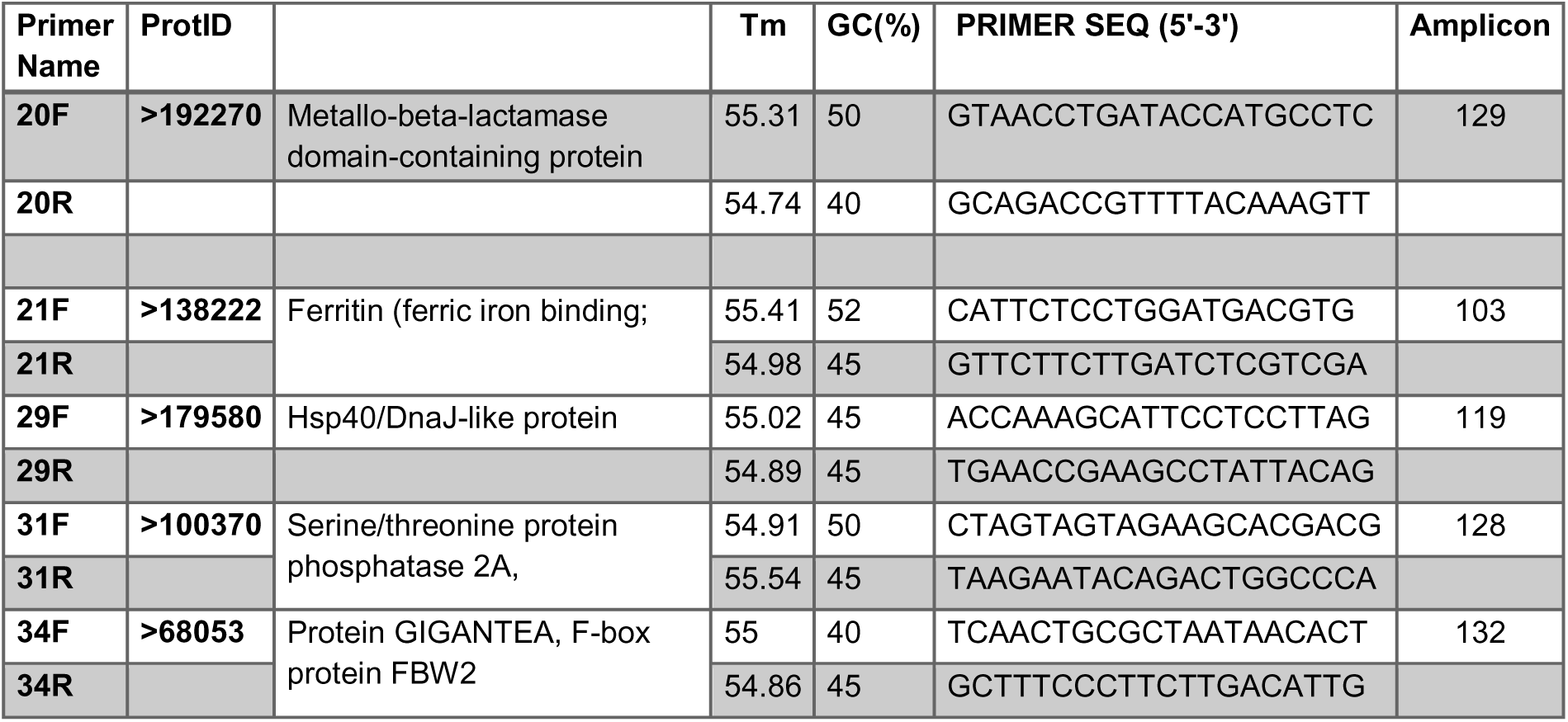
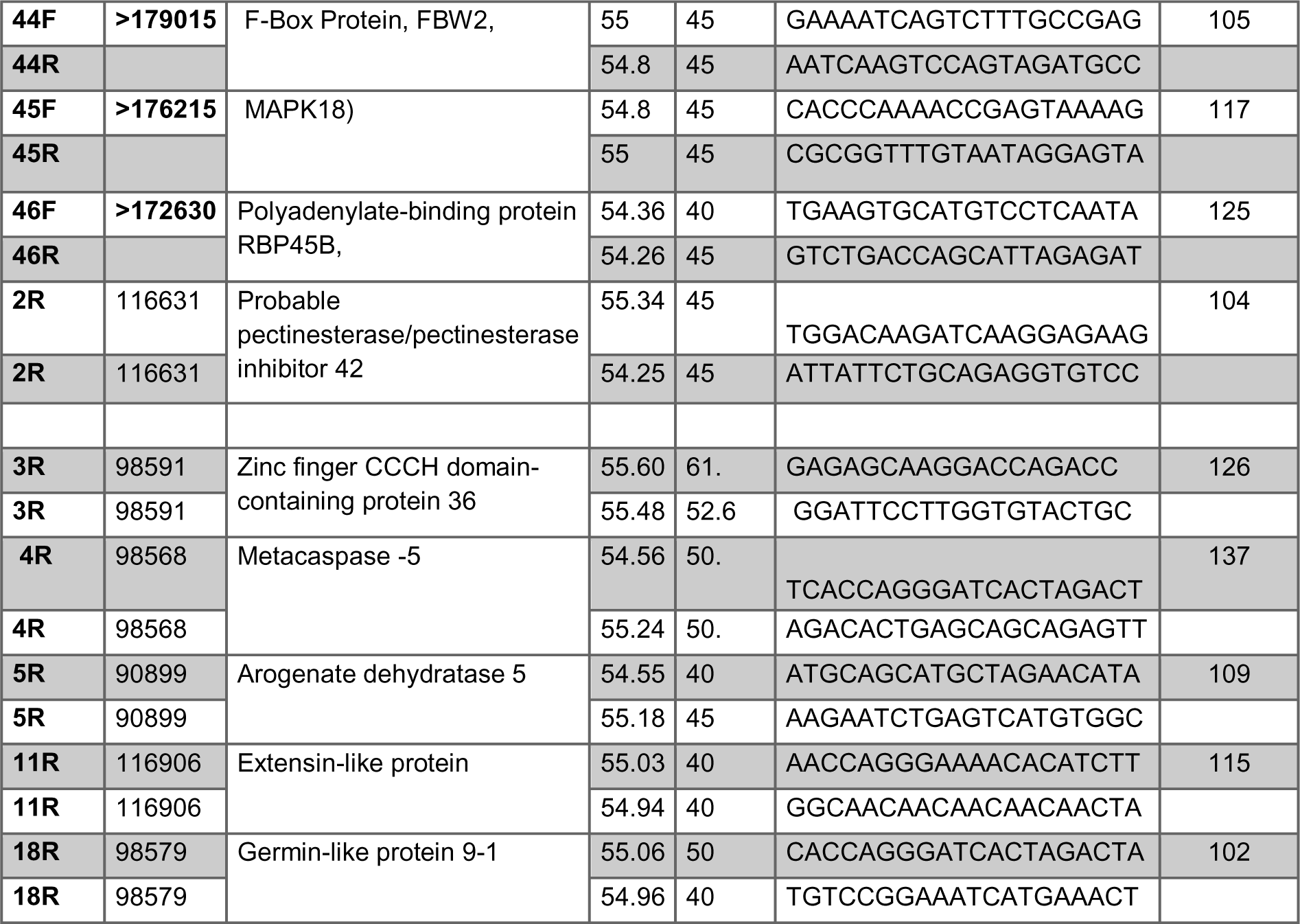
List of primers and their sequences used in the qRT-PCR experiments.

## REFERENCES

1. http://www.fao.org/worldfoodsituation/csdb/en/ (accessed on October 10, 2022).

2. Zampieri, M., Ceglar, A., Dentener, F., and Toreti, A. (2017). Wheat yield loss attributable to heat waves, drought and water excess at the global, national and subnational scales. Res. Lett. 12, 064008. 10.1088/1748-9326/aa723.

3. TUIK (2020), Crop production statistics, http://www.turkstat.gov.tr/ (accessed on September 14, 2020).

4. Manickavelu, A., Kawaura, K., Oishi, K., Shin, T., Kohara, Y., Yahiaoui, N., Keller, B., Abe, K., Suzuki, A., Nagayama, T., Yano, K., Ogihara, Y. (2012). Comprehensive Functional Analyses of Expressed Sequence Tags in Common Wheat (*Triticum aestivum L*.) DNA Res. 19, (2), 165–177. 10.1093/dnares/dss001.

5. Langridge, P., and Reynolds, M. (2021). Breeding for drought and heat tolerance in wheat Theoretical and Applied Genetics. 134, 1753–1769. 10.1007/s00122-021-03795-1.

6. Bartels, D., and Nelson, D. (1994). Approaches to improve stress tolerance using molecular genetics. *Plant*, Cell and Environ. 17, (5), 659–667. 10.1111/j.1365-3040.1994.tb00157.x.

7. Bohnert, H.J., Nelson, D.E., Jensen, R.G. (1995). Adaptations to Environmental Stresses. Plant Cell. 7, (7), 1099–1111. 10.1105/tpc.7.7.1099.

8. Mahajan, S., and Tuteja, N. (2005). Cold, Salinity and Drought Stresses: An Overview. Biochem Biophys. 444, 139–158. 10.1016/j.abb.2005.10.018.

9. del Pozo, J., and Ramirez-Parra, C.E. (2014). Deciphering the molecular bases for drought tolerance in Arabidopsis autotetraploids. Plant Cell Environ. 37, 2722–2737. 10.1111/pce.12344.

10. Iuchi, S., Kobayashi, M., Taji, T., Naramoto, M., Seki, M., Kato, T., Tabata, S., Kakubari, Y., Yamaguchi-Shinozaki, K., Shinozaki, K. (2001). Regulation of drought tolerance by gene manipulation of 9-cis-epoxycarotenoid dioxygenase, a key enzyme in abscisic acid biosynthesis in Arabidopsis. Plant J. 27, 325–333. 10.1046/j.1365-313x.2001.01096.x.

11. Fleury, D., Jefferies, S., Kuchel, H.; Langridge, P. (2010). Genetic and genomic tools to improve drought tolerance in wheat. J Exp Bot. 61, 3211–3222. 10.1093/jxb/erq152.

12. Kulik, A., Wawer, I., Krzywinska, E., Bucholc, M., Dobrowolska, G. (2011). SnRK2 protein kinases-Key regulators of plant response to abiotic stresses. OMICS. 15, 859–860. 10.1089/omi.2011.0091.

13. Movahedi, S., Sayed-Tabatabaei, B.E., Alizade, H., Ghobadi, C., Yamchi, A., Khaksar, G. (2012). Constitutive expression of Arabidopsis DREB1B in transgenic potato enhances drought and freezing tolerance. Biol Plant. 56, 37–42. 10.1007/s10535-012-0013-6.

14. Reguera, M., Peleg, Z., Blumwald, E. (2012). Targeting metabolic pathways for genetic engineering abiotic stress-tolerance in crops. Biochim Biophys Acta. 1819, 186–194. 10.1016/j.bbagrm.2011.08.005.

15. Mehrotra, R., Bhalothia, P., Bansal, P., Basantani, M.K., Bharti, V., Mehrotra, S. (2014). Abscisic acid and abiotic stress tolerance-Different tiers of regulation. J. Plant Physiol. 171, 486–496. 10.1016/j.jplph.2013.12.007.

16. Ding, L., Gao, C., Li, Y., Li, Y., Zhu, Y., Xu, G., Shen, Q., Kaldenhoff, R., Kai, L., Guo, S. (2015). The enhanced drought tolerance of rice plants under ammonium is related to aquaporin (AQP). Plant Sci. 234, 14–21. 10.1016/j.plantsci.2015.01.016

17. Liang, L., Wang W., Wang, J., Ma J., Li, C., Zhou, F., Zhang S., Yu Y., Zhang L., Li, W., Huang, X. (2017). Identifcation of diferentially expressed genes in sunfower (*Helianthus annuus*) leaves and roots under drought stress by RNA sequencing. Botanical Studies. 58, (42). 10.1186/s40529-017-0197-3.

18. Li, S., Pan Z., Chen, L., Dai Y., Wan J., Ye H., Nguyen, HT., Zhang G., and Chen, H. (2020) Analysis of Whole Transcriptome RNA-seq Data Reveals Many Alternative Splicing Events in Soybean Roots under Drought Stress Conditions. Genes. 11, 1520, 10.3390/genes 11121520.

19. Peremarti, A., Marè, C., Aprile, A., Roncaglia, E., Cattivelli, L., Villegas, D., et al.. (2014). Transcriptomic and proteomic analyses of a pale-green durum wheat mutant shows variations in photosystem components and metabolic deficiencies under drought stress. BMC Genomics. 15, 125. 10.1186/1471-2164-15-125.

20. Ma, J., Li, R., Wang, H., Li, D., Wang, X., Zhang, Y., et al.. (2017). Transcriptomics analyses reveal wheat responses to drought stress during reproductive stages under field conditions. Front. Plant Sci. 8, 592. 10.3389/fpls.2017.00592.

21. Cui, X., Wang, Y.-X., Liu, Z.-W., Wang, W.-L., Li, H., Zhuang, J. (2018). Transcriptome-wide identification and expression profile analysis of the bHLH family genes in Camellia sinensis. Funct Integr Genom. 18, 489–503. htpps://doi.org/10.1007/s10142-018-0608-x.

22. Close, T.J. (1996). Dehydrins: emergence of a biochemical role of a family of plant dehydration proteins. Physiol Plantarum. 97, (4), 795–803. doi.org/10.1111/j.1399-3054.1996.tb00546.x

23. Trouverie, J., TheÂvenot, C., Rocher, J-P., Sotta, B., and Prioul, J-L. (2003). The role of abscisic acid in the response of a specific vacuolar invertase to water stress in the adult maize leaf. J Exp Bot. 54, (390), 2177–2186. 10.1093/jxb/erg234.

24. Pnueli, L., Hallak-Herr, E., Rozenberg, M., Cohen, M., Goloubinoff, P., Kaplan, A., Mittler, R. (2002). Molecular and biochemical mechanisms associated with dormancy and drought tolerance in the desert legume *Retama raetam*. The Plant J. 31, (3), 319–330. 10.1046/j.1365-313X.2002.01364.x.

25. Anderson, J.V., Davis, D.G. (2004). Abiotic stress alters transcript profiles and activity of glutathione S-transferase, glutathione peroxidase, and glutathione reductase in Euphorbia esula. Physiol Planta.120, 421–433. 10.1111/j.0031-9317.2004.00249.x

26. Magwanga, R.O., Lu, P., Kirungu J.N., Lu H., Wang, X., Cai, X., Zhou, Z., Zhang Z., Salih., H., Wang K., Liu, F. (2018). Characterization of the late embryogenesis abundant (LEA) proteins family and their role in drought stress tolerance in upland cotton. BMC Genet. 19: 6. doi: 10.1186/s12863-017-0596-1

27. Cellier, F., Conéjéro, G., Breitler, J-C., Casse, F. (1998). Molecular and Physiological Responses to Water Deficit in Drought-Tolerant and Drought-Sensitive Lines of Sunflower: Accumulation of Dehydrin Transcripts Correlates with Tolerance. Plant Physiol. 116, (1), 319–328. 10.1104/pp.116.1.319.

28. Lopez, C.G., Banowetz, G.M., Peterson, C. J., Kronstad, W.E. (2003). Dehydrin Expression and Drought Tolerance in Seven Wheat Cultivars. Crop Science. 43, (2), 577–582. 10.2135/cropsci2003.5770.

29. Bogard, M., Hourcade, D., Piquemal, B., Gouache, D., Deswartes, J.-C., Throude, M., Cohan, J-P. (2021). Marker-based crop model-assisted i[]deotype design to improve avoidance of abiotic stress in bread wheat. J Exp Botany. 72, (4), 1085–1103. 10.1093/jxb/eraa477.

30. Barrs, H.D., and Weatherley, P.E. (1962). A Re-Examination of the Relative Turgidity Techniques for Estimating Water Deficits in Leaves. Aust J Biol Sci. 15, 413–428. 10.1071/BI9620413.

31. Keskin, B.C.., Topal-Sarikaya, A., Yuksel, B., Memon, A.R. (2010). Abscisic acid regulated gene expression in bread wheat (*Triticum aestivum* L.). Aust J Crop Science 4, 617–625. 10.3316/informit.857732547077080.

32. Andrews S. FastQC: a quality control tool for high throughput sequence data. Available online at http://www.bioinformatics.babraham.ac.uk/projects/fastqc. Accessed 15 Oct 2022.

33. FASTX-Toolkit: FASTQ/a short-reads pre-processing tools. http://hannonlab.cshl.edu/fastx_toolkit/. Accessed Jan 2022.

34. Duan, J., Xia, C., Zhao, G., Jia, J., Kong, X. (2012). Optimizing de novo common wheat transcriptome assembly using short-read RNA-Seq data. BMC Genomics. 13, 392, 1–12. 10.1186/1471-2164-13-392.

35. Robinson, M.D., McCarthy, D.J., Smyth, G.K. (2010). edgeR: a Bioconductor package for differential expression analysis of digital gene expression data. Bioinformatics. 26, (1), 139–140. 10.1093/bioinformatics/btp616.

36. https://rnabio.org/module-07-trinotate/0007/02/01/Trinotate/

37. Cevher-Keskin, B., Yuca, E., Ertekin, O., Yuksel, B., Memon, A.R. (2011). Expression characteristics of ARF1 and SAR1 during development and the de-etiolation process. Plant Biol. htpps://doi:10.1111/j.1438-8677.2011.00482.x.

38. Schmittgen, T.D., Zakrajsek, B.A., Mills, A.G., Gorn, V., Singer, M.J., Reed, M.W. (2000). Quantitative Reverse Transcription–Polymerase Chain Reaction to Study mRNA Decay: Comparison of Endpoint and Real-Time Methods. Analytical Biochem. 285, (2), 194–204. 10.1006/abio.2000.4753.

39. http://bioinfo.cau.edu.cn/agriGO/

40. Kohli, P., Kalia, M., Gupta, R. (2015). Pectin Methylesterases. J Bioprocess Biotech. 5, 3–7. 10.4172/2155-9821.1000227.

41. Cassab, G. (1998). Plant cell wall proteins. Annu Rev Plant Physiol Plant Mol Biol. 49, 281–309. 10.1146/annurev.arplant.49.1.281.

42. Bernier, F., Berna, A. (2001). Germins and germin-like proteins: plant do-all proteins. But what do they do exactly? Plant Physiol Biochem. 39, 545–554. doi:10.1016/s0981-9428(01)01285-2.

43. Lou, Y., and Baldwin, I.T. (2006). Silencing of a germin-like gene in *Nicotiana attenuata* improves performance of native herbivores. Plant Physiol. 140 (3), 1126–1136. 10.1104/pp.105.073700.

44. Valandro, F., Menguer, P.K., Cabreira-Cagliari, C., Margis-Pinheiro, M., Cagliari, A. 2020. Programmed cell death (PCD) control in plants: New insights from the Arabidopsis thaliana deathosome. Plant Science. 299, 110603. 10.1016/j.plantsci.2020.110603.

45. van der Oost, J., Huynen, M.A., Verhees, C.H. (2002). Molecular characterization of phosphoglycerate mutase in archaea FEMS. Microbiology Letters. 212, (1) 111–120. 10.1111/j.1574-6968.2002.tb11253.x.

46. Kataya, A.R.A., Heidari, B., Hagen, L., Kommedal, R., Slupphaug, G., Lillo, C. (2015). Protein phosphatase 2A holoenzyme is targeted to peroxisomes by piggybacking and positively affects peroxisomal β-oxidation. Plant Physiol. 167 (2), 493–506. 10.1104/pp.114.254409.

47. Fowler, S., Lee, K, Onouchi, H., Samach, A., Richardson, K., Morris, B., Coupland, G., Putterill, J. (1999). GIGANTEA: a circadian clock-controlled gene that regulates photoperiodic flowering in Arabidopsis and encodes a protein with several possible membrane-spanning domains The EMBO J. 18, 4679–4688. 10.1093/emboj/18.17.4679.

48. Abdel-Ghany, S.E., Ullah, F., Ben-Hur, A., Reddy, A.S.N. (2020). Transcriptome Analysis of Drought-Resistant and Drought-Sensitive Sorghum (*Sorghum bicolor*) Genotypes in Response to PEG-Induced Drought Stress. Int. J Mol Sci. 21, (772). 10.3390/ijms 21030772.

49. Wang, X., Bao, K., Reddy U.K., Bai, Y., Hammar, S. A., Jiao, C., Wehner, T.C., Ramírez-Madera, A.O., Weng, Y., Grumet, R., Fei, Z. (2018). The USDA cucumber (*Cucumis sativus* L.) collection: genetic diversity, population structure, genome-wide association studies, and core collection development. Horticulture Research. 5, (64), 10.1038/s41438-018-0080-8.

50. Moumeni A., Satoh K., Kondoh H, Asano T., Hosaka A., Venuprasad R., Serraj R., Kumar A., Leung H., Kikuchi S. (2011). Comparative analysis of root transcriptome profiles of two pairs of drought-tolerant and susceptible rice near-isogenic lines under different drought stress. BMC Plant Biol. 2011, 11, (174), 1-17. http://www.biomedcentral.com/1471-2229/11/174.

51. Pape, H.-C., and Pare, D. (2010). Plastic Synaptic Networks of the Amygdala for the Acquisition, Expression, and Extinction of Conditioned Fear. Physiol. Rev. 90, 419–463. 10.1152/physrev.00037.2009.

52. Wang, D., Guo, Y., Wu, C., Yang, G., Li, Y., Zheng, C.. (2008). Genome-wide analysis of CCCH zinc finger family in Arabidopsis and rice. BMC Genomics. 9, 44. 10.1186/1471-2164-9-44.

53. Pomeranz, M.C., Hah, C., Lin, P.C., Kang, S.G., Finer, J.J., Blackshear, P.J., et al. (2010). The Arabidopsis tandem zinc finger protein AtTZF1 traffics between the nucleus and cytoplasmic foci and binds both DNA and RNA. Plant Physiol. 152, 151–165. 10.1104/pp.109.145656

54. Anderson, P., and Kedersha, N. (2009). RNA granules. Nat Rev Mol Cell Biol. 10, 430–436. 10.1038/nrm2694.

55. Lin, P.C., Pomeranz, M.C., Jikumaru, Y., Kang, S.G., Hah, C., Fujioka, S., Kamiya, Y., Jang, J.C. (2011). The Arabidopsis tandem zinc finger protein *AtTZF1* affects ABA- and GA-mediated growth, stress and gene expression responses. The Plant J. 65, 253–268. 10.1111/j.1365-313X.2010.04419.x

56. Lee, S.J., Jung, H.J., Kang, H., Kim, S.Y. (2012). Arabidopsis zinc finger proteins AtC3H49/AtTZF3 and AtC3H20/AtTZF2 are involved in ABA and JA responses. Plant Cell Physiol. 53, 673–686. 10.1093/pcp/pcs023.

57. Balla, G., Jacob, H.S., Balla, J., Rosenberg, M., Nath, K., Apple, F., Eaton, J.W., Vercellotti, G.M. (1992). Ferritin: a cytoprotective antioxidant strategem of endothelium. J Biol Chem. 267, 18148–18153. 10.1016/S0021-9258(19)37165-0.

58. Fobis-Loisy, I., Loridon, K., Lobreaux, S., Lebrun, M., Briat, J.F. (1995). Structure and differential expression of two maize ferritin genes in response to iron and abscisic acid. Eur J Biochem. 231, 609–619. 10.1111/j.1432-1033.1995.0609d.x

59. Van Wuytswinkel, O., Briat, J.F. (1995). Conformational changes and *in vitro* core formation modifications induced by site directed mutagenesis of the specific amino terminus (EP) of pea seed ferritin. The Biochem J. 305, 959–965. 10.1042/bj3050959.

60. Murgia, I., Delledonne, M., Soave, C. (2002). Nitric oxide mediates iron-induced ferritin accumulation in Arabidopsis. The Plant J. 30, (5), 521–528. 10.1046/j.1365-313X.2002.01312.x

61. Al-Qsous, S., Carpentier, E., Klein-Eude, D., Burel, C., Mareck, A., Dauchel, H., Gomord, V., Balangé, A.P. (2004). Identification and isolation of a pectin methylesterase isoform that could be involved in flax cell wall stiffening. Planta. 219, 369–378. 10.1007/s00425-004-1246-1.

62. Micheli, F. (2001). Pectin methylesterases: cell wall enzymes with important roles in plant physiology. Trends Plant Sci. 6, 414–419. 10.1016/S1360-1385.

63. Cramer, G.R., Ergül, A., Grimplet, J., Tillett, R. L., Tattersall, E.A.R., Bohlman, M.C., Vincent, D., Sonderegger, J., Evans, J., Osborne, C., Quilici, D., Schlauch, K.A., Schooley, D. A., Cushman, J. C. (2007). Water and salinity stress in grapevines: early and late changes in transcript and metabolite profiles. Funct Integr Genom. 7, 111–134. htpps://doi.org/10.1007/s10142-006-0039-y.

64. Houde, M., and Diallo, A.O. (2008). Identification of genes and pathways associated with aluminum stress and tolerance using transcriptome profiling of wheat near-isogenic lines BMC Genomics. 9, 400, 1–13. htpps://doi.org/10.1186/1471-2164-9-400.

65. Ke, Y., Han, G., He, H., Li, J. (2009). Differential regulation of proteins and phosphoproteins in rice under drought stress. Biochem Biophys Res Commun. 379, (1), 133–138. 10.1016/j.bbrc.2008.12.067.

66. Cevher-Keskin, B. (2019). Quantitative mRNA expression profiles of germin-like and extensin-like proteins under drought stress in *Triticum aestivum* L. Int J Life Sci Biotech. 2 (2), 95–107. 10.38001/ijlsb.566942.

67. Ficke, A., Gadoury, D.M., Seem, R.C. (2002). Ontogenic resistance and plant disease management: a case study of grape powdery mildew. Phytopath. 92 (6), 671–675. 10.1094/PHYTO.2002.92.6.671.

68. Manosalva, P., Davidson, R., Liu, B., Zhu, X., Hulbert, S., Leung, H., Leach, J. (2009). A germin-like protein gene family functions as a complex qtl conferring broad-spectrum disease resistance in rice. Plant Physiol. 149 (1), 286–296. 10.1104/pp.108.128348.

69. Knecht, K., Seyffarth, M., Desel, C., Thurau, T., Sherameti, I., Lou, B., Oelmüller, R., Cai, D. (2010). Expression of bvglp-1 encoding a germin-like protein from sugar beet in *Arabidopsis thaliana* leads to resistance against phytopathogenic fungi. Mol Plant Microb Interact. 23, (4), 446–457. 10.1094/MPMI-23-4-0446.

70. Christensen, A., Thordal-Christensen, H., Zimmermann, G., Gjetting, T., Lyngkjaer, M., Dudler, R., Schweizer, P. (2004). The germin like protein GLP4 exhibits superoxide dismutase activity and is an important component of quantitative resistance in wheat and barley. Mol Plant Microbe Interact. 17, 109–117. 10.1094/MPMI.2004.17.1.109.

71. Corea, O.R.A., Ki, C., Cardenas, C.L., Kim, S.-J., Brewer, S.E., Patten, A.M., Davin, L.B., Lewis, N.G. (2012a). Arogenate Dehydratase Isoenzymes Profoundly and Differentially Modulate Carbon Flux into Lignins. J Biol Chem. 287, (14), 11446–11459. 10.1074/jbc.M111.322164.

72. Corea, O.R.A., Bedgar, D.L., Davin, L.B., Lewis, N.G. (2012b). The arogenate dehydratase gene family: towards understanding differential regulation of carbon flux through phenylalanine into primary versus secondary metabolic pathways. Phytochem. 82, 22–37. htpps://doi.org/10.1016/j.phytochem.2012.05.026

73. Chen, Q., Man, C., Li, D., Tan, H., Xie, Y., Huang, J. (2016). Arogenate Dehydratase Isoforms Differentially Regulate Anthocyanin Biosynthesis in *Arabidopsis thaliana*. Mol Plant. 10.1016/j.molp.2016.09.010.

74. Uren, A.G., O’Rourke, K., Aravind, L., Pisabarro, M.T., Seshagiri, S., Koonin, E.V., Dixit V.M. (2000). Identification of paracaspases and metacaspases: two ancient families of caspase-like proteins, one of which plays a key role in MALT lymphoma Mol Cell. 6, (4), 961–967.

75. Huh, S.U. (2022) Evolutionary Diversity and Function of Metacaspases in Plants: Similar to but Not Caspases. Int J Mol Sci. 223, (9), 4588. 10.3390/ijms23094588.

76. Oslund, R.C., Su, X., Haugbro, M., Kee, J-M., Esposito, M., David, Y., Wang, B., Ge, E., Perlman, D.H., Kang, Y., Muir, T.W., Rabinowitz, J.D. (2017). Bisphosphoglycerate mutase controls serine pathway flux via 3- phosphoglycerate. Nat Chem Biol. 13, (10), 1081–1087. 10.1038/nchembio.2453.

77. Chen, J., Zhu, X., Shen, G., Zhang, H. (2015). Overexpression of *AtPTPA* in Arabidopsis increases protein phosphatase 2A activity by promoting holoenzyme formation and ABA negatively affects holoenzyme formation. Plant Signal Behav. 10, (11), e1052926. htpps://doi: 10.1080/15592324.2015.1052926.

78. Creighton, M.T., Sanmartín, M., Kataya, A.R.A., Averkina, I.O., Heidari, B., Nemie-Feyissa, D., Sánchez-Serrano, J.J., Lillo, C. (2017). Light regulation of nitrate reductase by catalytic subunits of protein phosphatase 2A. Planta. 246 (4), 701–710. htpps://doi.org/ 10.1007/s00425-017-2726-4.

79. Yu, R.M.K., Zhou, Y., Xu, Z.F., Chye, M.L., Kong, R.Y.C. (2003). Two genes encoding protein phosphatase 2A catalytic subunits are differentially expressed in rice. Plant Mol Biol. 51, 295–311. 10.1023/a:1022006023273.

80. Pais, M., Tellez, M., Capiati D. (2009). Serine/Threonine Protein Phosphatases type 2A and their roles in stress signaling. Plant Signaling Behav. 4, (11), 1013–1015. DOI:10.4161/psb.4.11.9783.

81. Xu, S.M., Wang, X.C., Chen, C. (2007). Zinc finger protein1 (ThZF1) from salt cress (*Thellungiella halophila*) is a Cys-2/His-2-type transcription factor involved in drought and salt stress. Plant Cell Rep. 26, 497–506. 10.1007/s00299-006-0248-9.

82. Krahmer, J., Ganpudi, A., Abbas, A., Romanowski, A., Halliday, K. J. (2018). Phytochrome, Carbon Sensing, Metabolism, and Plant Growth Plasticity. Plant Physiol. 176, (2), 1039– 1048. 10.1104/pp.17.01437.

83. Waters, M.T., Scaffidi, A., Sun, Y.K., Flematti, G.R., Smith, S.M. (2014). The karrikin response system of Arabidopsis. The Plant J. 79, 623–631., 10.1111/tpj.12430.

84. Kim, W.-Y., Ali, Z., Park, H.J., Park, S.J., Cha, J.-Y., Perez-Hormaeche, J., Quintero, F.J., Shin, G., Kim, M.R., Qiang, Z., et al. (2013). Release of SOS2 kinase from sequestrationwith GIGANTEA determines salt tolerance in Arabidopsis. Nat Commun. 4, 1352. 10.1038/ncomms2357.

85. Riboni, M., Galbiati, M., onelli, C., Conti L. (2013). GIGANTEA Enables Drought Escape Response via Abscisic Acid-Dependent Activation of the Florigens and SUPPRESSOR OF OVEREXPRESSION OF CONSTANS1. Plant Physiol. 162, 3, 1706–1719. 10.1104/pp.113.217729.

86. Cao, S., Ye, M., Jiang, S. (2005). Involvement of GIGANTEA gene in the regulation of the cold stress response in Arabidopsis. Plant Cell Rep. 24, 683–690. 10.1007/s00299-005-0061.

87. Fornara, F., De Montaigu, A., Sanchez-Villarreal, A., Takahashi, Y., Van Themaat, E.V.L., Huettel, B., Davis, S.J., Coupland, G. (2015). The GI-CDF module ofArabidopsis affects freezing tolerance and growth aswell as flowering. Plant J. 81, 5, 695–706. 10.1111/tpj.12759.

88. Yan, Y., Gan J., Tao, Y., Okita, T.W., Tian L. (2022). RNA-Binding Proteins: The Key Modulator in Stress Granule Formation and Abiotic Stress Response. Front. Plant Sci. 13, 882596. 10.3389/fpls.2022.882596.

