## Supplementary figures and images for "Deciphering Drought Response Mechanisms: Transcriptomic Insights from Drought-Tolerant and Drought-Sensitive Wheat (*Triticum aestivum* L.) Cultivars"

### Supplemental Figure S1

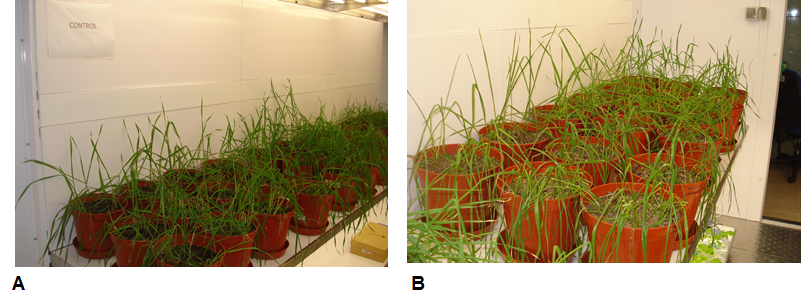

### Supplemental Figure S2

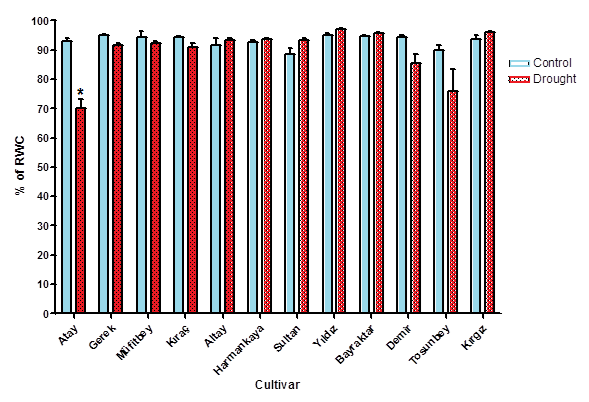

### Supplemental Figure S3

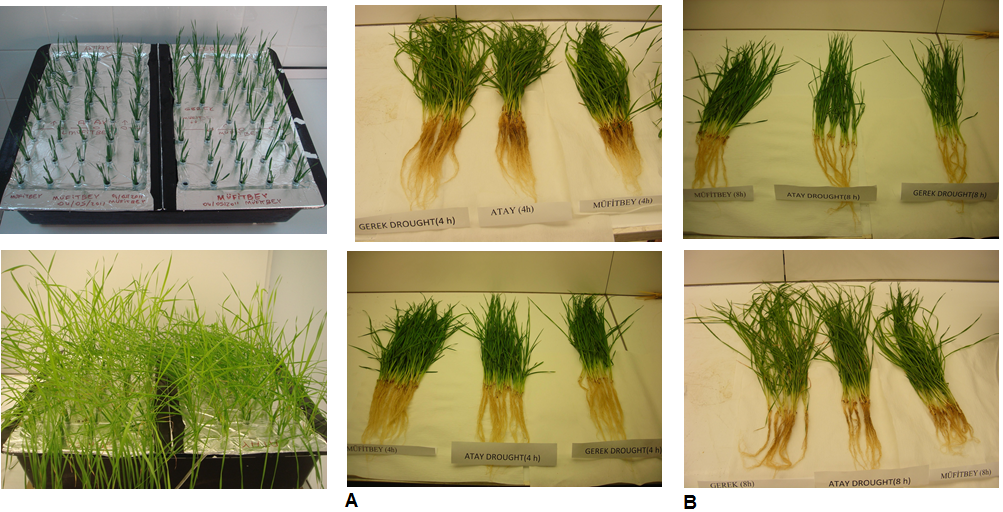

### Supplemental Figure S4

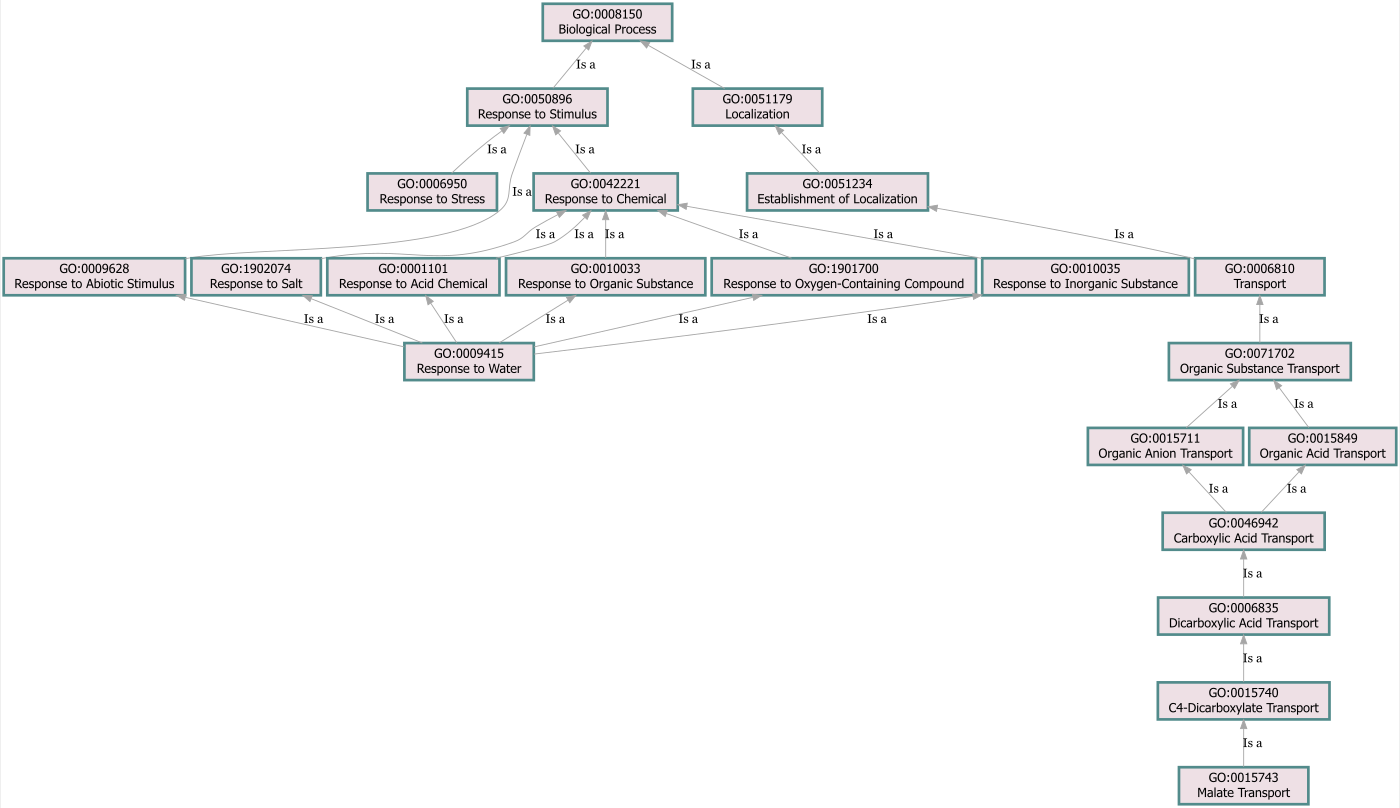

### Supplemental Figure S5

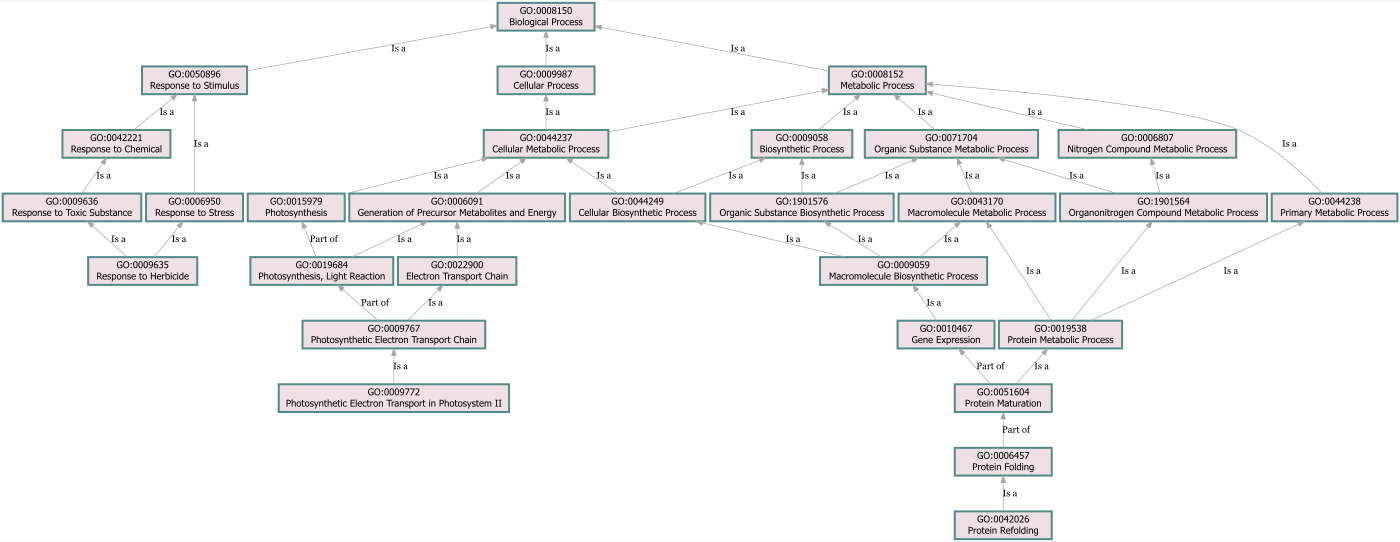

### Supplemental Figure S6

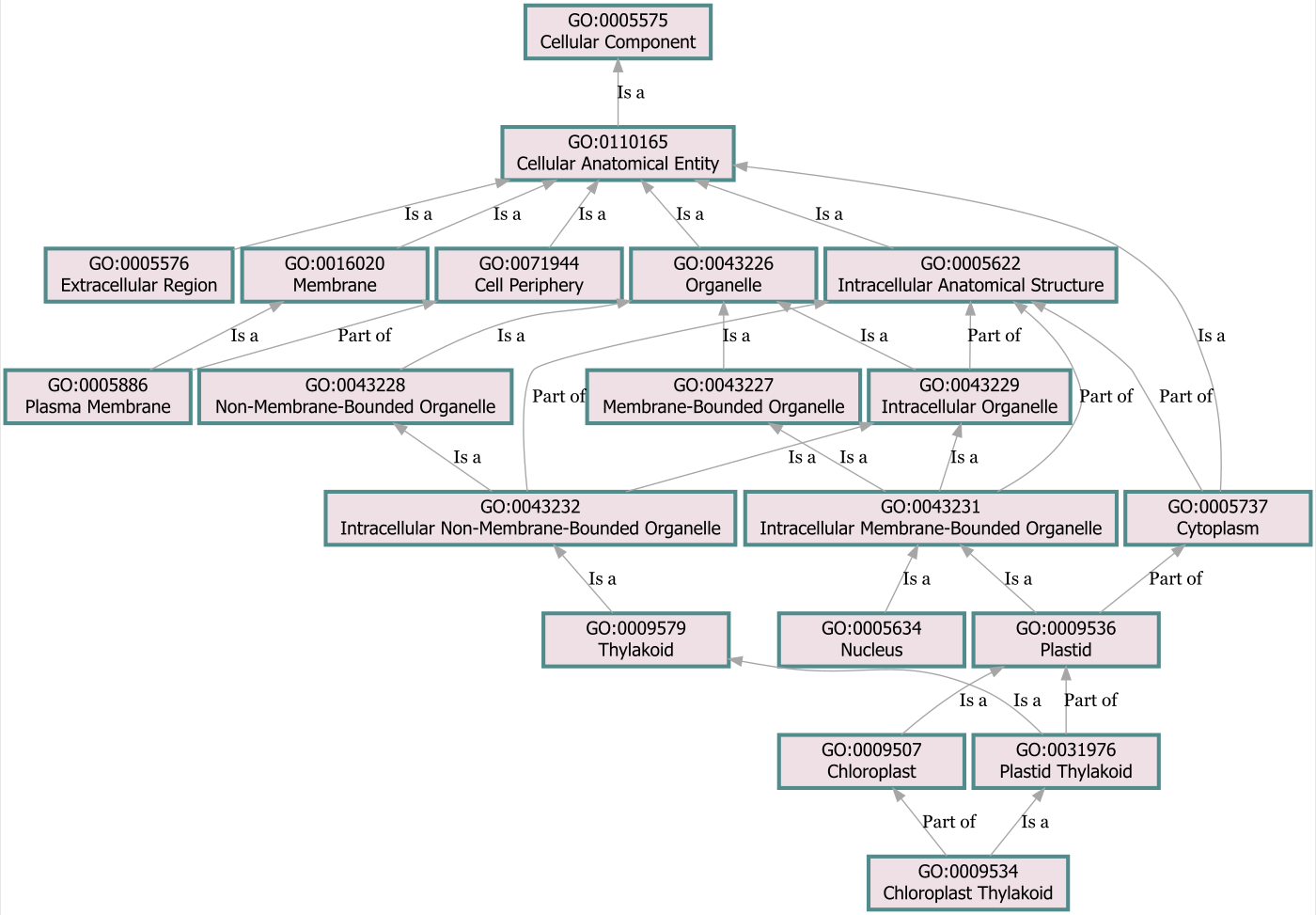

### Supplemental Figure S7

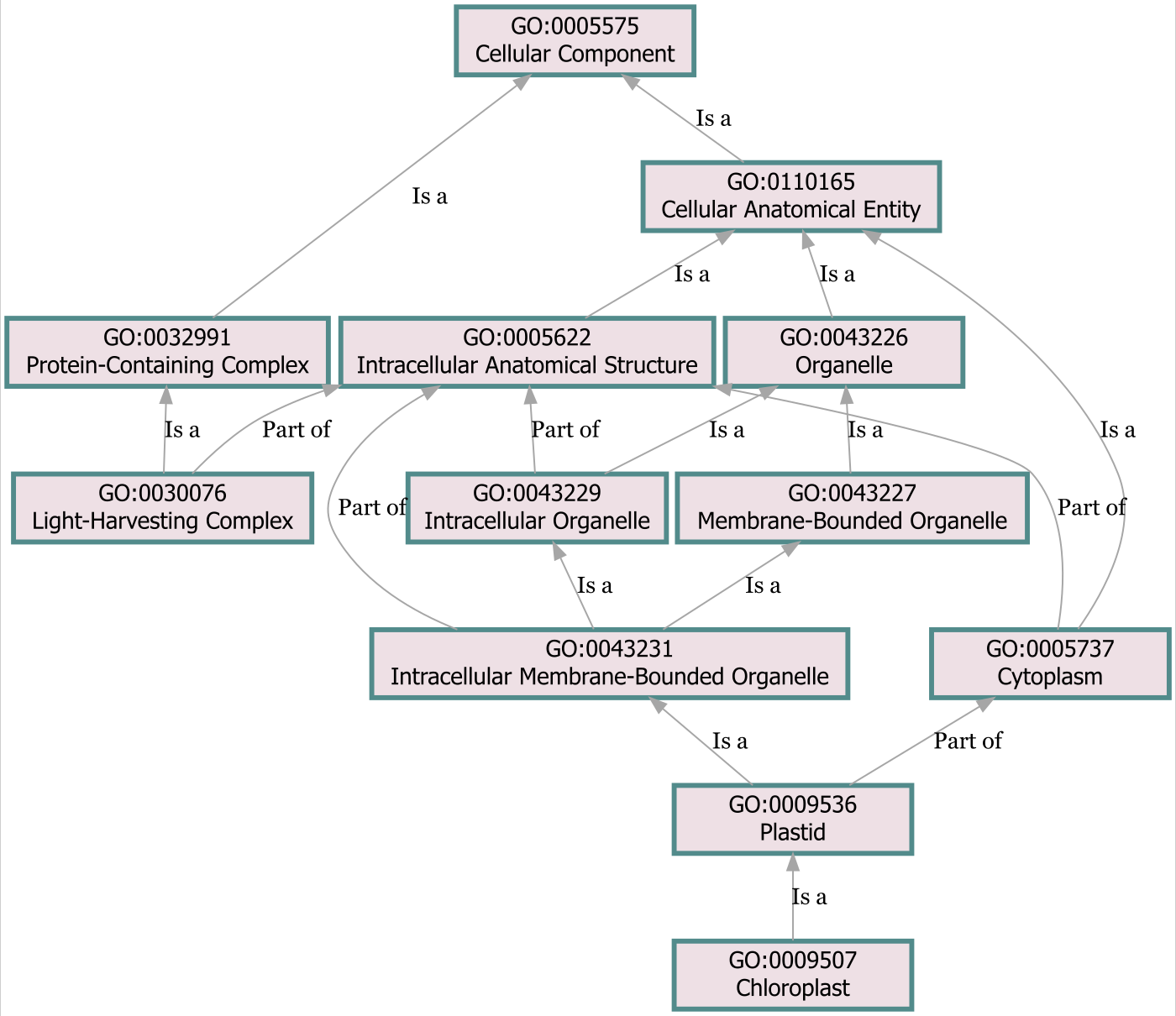

### Supplemental Figure S8

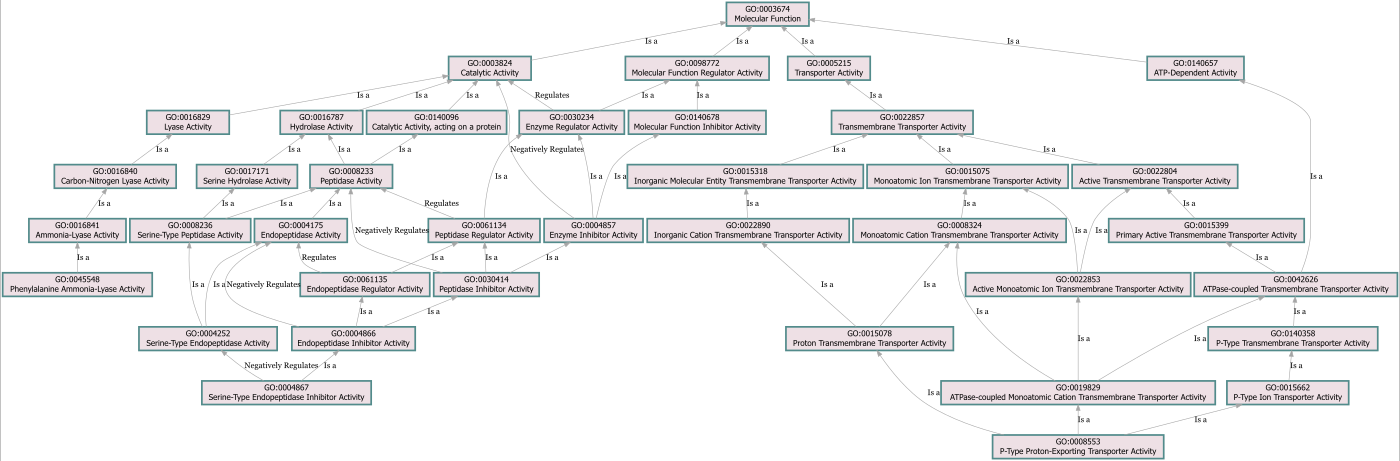

### Supplemental Figure S9

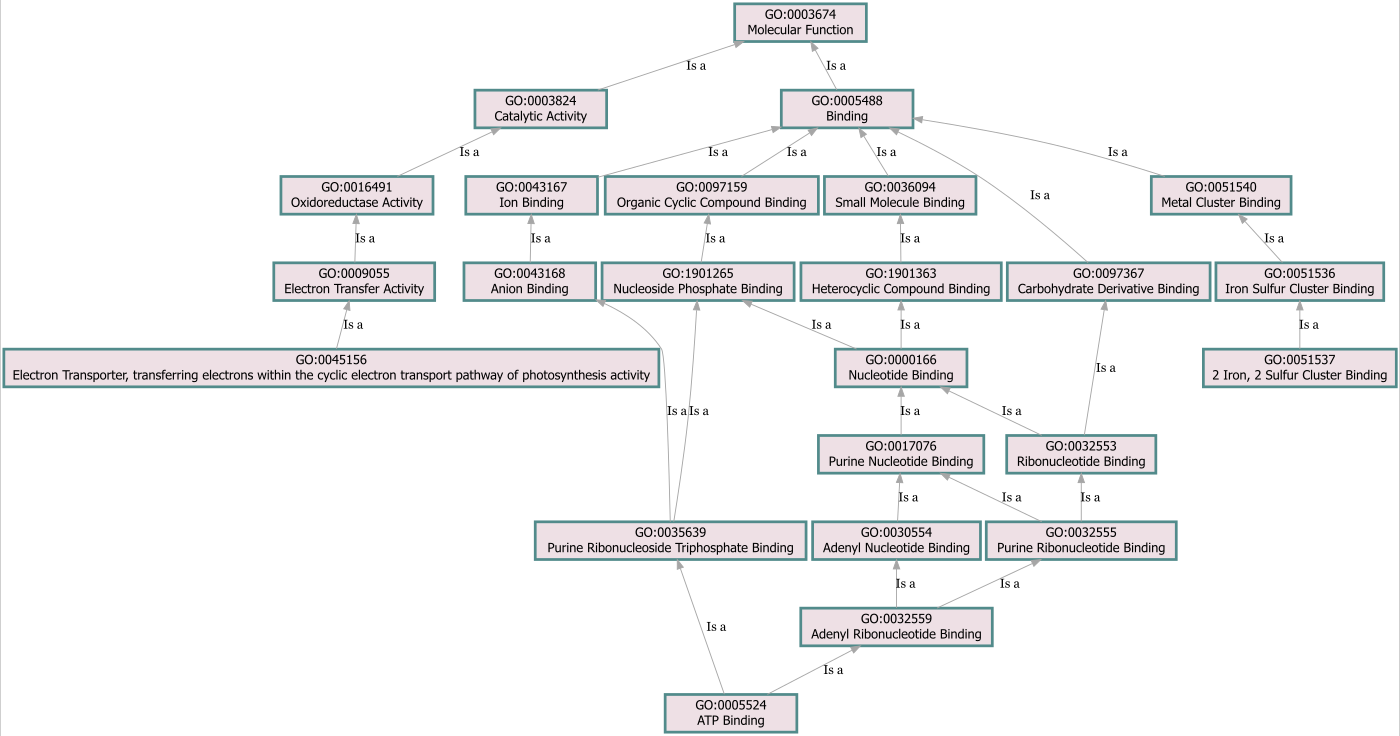

### Supplemental Figure S10

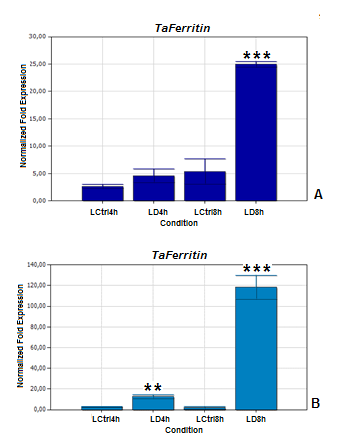

### Supplemental Figure S11

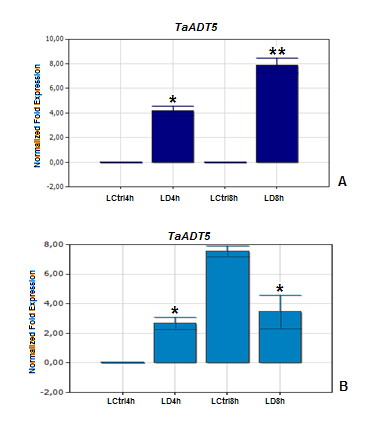
